# Patient-specific logical models replicate phenotype responses to psoriatic and anti-psoriatic stimuli

**DOI:** 10.1101/2023.08.24.554583

**Authors:** Eirini Tsirvouli, Eir Aker, Martin Kuiper

## Abstract

Psoriasis is a dermatologic disease that affects 2% of the world population. Psoriasis is characterized by chronic inflammation and aberrant behavior of keratinocytes, which display increased levels of proliferation, and decreased differentiation and apoptosis. Stimulation of keratinocytes by psoriatic cytokines leads to the increased production of immunostimulatory ligands that further attract immune cells and amplify inflammatory responses. Psoriasis can have severe, moderate, or mild outcomes and while these severity levels demand custom medical treatment schemes, assigning an effective treatment to patients with moderate or severe disease is a demanding task.

The varied responses of patients to treatments highlight a large disease complexity, demanding that new ways to analyze and integrate patients’ molecular profiles are developed to design patient-specific therapies. We have used gene expression values from psoriasis biopsies to separate patients into two clusters, each with distinct expression profiles, but nevertheless not correlating with any of the available clinical data, such as disease severity. When using these gene expression levels in logical model simulations these data became highly descriptive of patient-specific phenotype characteristics. Starting from a psoriatic keratinocyte model that we published recently, we added additional pathways highlighted by a differential gene expression analysis between the subgroups. This included components from the Interleukin-1 family, IFN-alpha/beta and IL-6 signaling pathways. Model personalization was performed by using patient gene expression levels in model configurations, exploiting the PROFILE pipeline. Personalized simulations revealed that the two patient clusters represent more innate immunity-driven, highly inflammatory phenotypes and adaptive immunity-driven, chronic phenotypes, respectively. The model was also able to finely capture differences between responses in patients with a known disease severity. A treatment response analysis among the patient cohort predicted differential responses to the inhibition of psoriatic stimuli, with IL-17, TNFα and PGE2 inhibition reducing proliferation and inflammatory phenotypes. Alternative treatment with PGE2 or TNFα inhibition instead of IL-17 was suggested for patients with high NF-κB activity and prosurvival factors, such as CREB1. With this project, we aim to highlight the value of combining omics data with logical modeling for the detection of ‘emergent’ phenotypes and for gaining disease knowledge on the individual patient level.

## 1. Introduction

Chronic inflammation is a hallmark of psoriasis: a dermatologic disease that affects two percent of the world population. Patients can display a spectrum of different phenotypic variants (psoriatic subtypes) but psoriasis vulgaris (or plaque psoriasis) is by far the most common psoriatic subtype affecting 90% of psoriasis patients. Phenotypically, plaque psoriasis manifests itself as skin eruptions consisting of defined skin thickening and regions of red erythema underneath layers of scaling.

Keratinocytes (KCs) play a key role in psoriasis, together with several cell types of the immune system that are showing dysregulation of processes similar to an autoimmune condition. Skin eruption is proposed to be related to the reactions of the immune system to triggers from environmental stimuli like physical and mental distress, dermatological insults, or bacterial infections (Zhang, 2019), or to genetic features which predispose individuals for psoriasis (Ran et al., 2019). KCs triggered by such internal or external factors can recruit plasmacytoid dendritic cells (pDCs) so that together they produce psoriasis-inducing cytokines (most typically Type I Interferons (IFN)), chemokines, and other chemoattractant molecules, in addition to antimicrobial and immunostimulatory peptides. One example of such peptides is Cathelicidin Antimicrobial Peptide (LL-37) (Albanesi et al., 2018), which, in concert with fragments of DNA/RNA from affected tissue or pathogens, can be recognized by receptors of DCs and commence an inflammatory cascade that will ultimately lead to psoriasis. The activation of receptors such as TLRs (Toll-like receptors) in pDCs mediates production of interleukins (mainly IL-23 and IL-12) and tumor necrosis factor alpha (TNFα). These cytokines then shape the differentiation of T-helper cells towards the Th17 and Th1 cell phenotypes. IL-17, Type 2 IFNs and IL-22 are important cytokines secreted by the aforementioned T-cells and have principal functions in the disease progression. KCs are able to respond to all these cytokines by altering their normal life cycle and by further secreting cytokines and chemokines, that further amplify immune responses and immune cell recruitment to the psoriatic lesions (Albanesi et al., 2018; Albanesi, 2019). These events establish a vicious cycle between immune cells and KCs, ultimately leading to a chronic inflammatory disease.

Psoriasis currently remains incurable, and due to its chronic nature, a lifelong treatment plan is needed for each patient. Psoriasis can be severe, moderate, or mild; each requiring specific medical treatment regimens. For mild psoriasis cases, the therapeutic strategy often consists of using creams which give topical relief. Generally, for mild psoriasis local treatment with corticosteroids or Vitamin D analogs have shown high effectiveness in alleviating disease symptoms (Armstrong and Read, 2020). In moderate-to-severe or severe cases there is a need for more systemic therapeutic options.

There, a new generation of drugs called biologics, have been used to treat psoriasis with unprecedented efficacy. Biologics predominantly target the activation of main components of the inflammatory axis that drives psoriasis pathogenesis, namely the TNFα/IL-23/IL-17 axis. As a result, the (hyper)activation of immune cells, such as dendritic cells and T-helper (Th) 17 cells can be inhibited. However, as the same cell types play a significant role in inflammatory and immune cell responses, treatment with biologics can be accompanied by several side effects, such as recurrent infections. Additionally, there is a significant proportion of patients who remain either non-responsive to current therapies or experience relapse of the disease (Armstrong and Read, 2020).

The identification of disease biomarkers is a critical enabling part of precision medicine. Such markers can inform about the risk of developing a disease, its progression or severity, and response to therapy. For psoriasis, more than 80 susceptibility genes and loci have been identified (Ran et al., 2019). Most of the susceptibility genes are related to skin barrier functions (e.g., Late cornified envelope (LCE), which plays a role in epidermal terminal differentiation), innate immune responses (e.g., TNFAIP3 and TNIP1 genes, which regulate NF-κB signaling) and adaptive immune responses (e.g., variants of the IL-23 receptor, which regulates Th-17 differentiation and expansion) (Ran et al., 2019). Other aspects of the disease, such as onset age and severity, have been linked to specific biomarkers (Ramessur et al., 2022). IL-36 is an example of such a biomarker as its expression was found to be significantly higher in critical cases such as generalized pustular psoriasis and high degree psoriasis vulgaris. The effects of IL-36 cytokines may be locally confined to skin lesions, but can also become a systemic issue which can aggravate disease symptoms (Catapano et al., 2020). Further studies suggest that serum levels of the interleukins IL-17 and IL-22 have an association to disease severity (Michalak-Stoma et al., 2013). Lastly, also a few biomarkers have been identified that inform about the response to systemic treatment (Corbett et al., 2022).

In this project, we aimed to group patients based on their global gene expression profiles and enhance the psoriatic keratinocyte (PsoKC) logical model developed previously (Tsirvouli et al., 2021) to better represent these patients and allow analysis of regulatory pathways underlying these expression differences. The PsoKC model represents the behavior of psoriatic KCs upon the activation of cytokine and Prostaglandin E2 (PGE2) signaling. This PsoKC model was extended with the biological pathways showing the highest variation across the patients and documented in literature to be important for psoriasis. These pathways were identified by analyzing public gene expression data from 185 psoriatic patients, which were classified into two gene expression clusters. Interestingly, these two clusters did not separate patients for disease severity or other available clinical variables, despite fairly conspicuous gene expression differences between these groups. The most differentially expressed regulatory pathways with psoriasis-significance were added to the PsoKC model and this extended model was used as a tool to develop patient-specific models and identify functional differences between the two patient clusters, using the expression profiles of individual patients to reveal differential disease phenotypes between the two patient clusters and treatment responses.

## 2. Materials & Methods

### 2.1 Transcriptomics data sets

In this project we used public gene expression data from psoriatic patients, collected, homogenized, and analyzed previously (Federico et al., 2020). As these data includes several datasets of different properties which could result in noise for the patient subgroup identification, only a selection of the available datasets were used further. The selection was based on several criteria, including disease, sample origin, treatment status, and detection method; selecting only RNA-Sequencing (RNA-seq) transcriptomics from samples originating from untreated lesional or non-lesional skin. Additionally, datasets from healthy donor skin samples were included as control. Based on the above selection criteria, nine datasets, comprising 370 unique patient samples, qualified for further analysis. An overview of the selected datasets can be found in Supplementary Table 1.

#### 2.1.1 Data pre-processing

The RNA-seq count matrices were processed by Federico et al. in a consistent manner, namely: filtering of poorly expressed RNA transcripts and upper quantile normalization (Federico et al., 2020). The nine selected datasets were combined into one expression data frame with a total of 370 individual samples and 12.779 genes. Log2 transformation was carried out to make the variance range in the expression data homoskedastic, and to reduce the substantial influence of high count genes. To further reduce experimental and technical dissimilarities between the individual studies, batch effect correction was performed with the RemoveBatchEffects function of the limma R package (Ritchie et al., 2015). A principal component analysis (PCA) was carried out to validate the batch correction and ensure that samples were separated by tissue of origin (psoriatic lesion, non-involved skin or healthy controls). These analyses were performed with PCAtools version 2.4.0 (Blighe et al., 2022). All data processing and analyses have been performed with R (R Core Team, 2013) in R studio version 1.4.1717.

#### 2.1.2 Differential expression analysis

Differential gene expression analysis was carried out by comparing the psoriatic patient samples with the healthy donors (baseline samples). Version 3.34.1 of EdgeR (Robinson et al., 2010) was employed, with the raw expression values of the combined datasets as input. Genes with no or only a low expression were removed and Trimmed Mean of M-values (TMM) normalization was conducted. Differentially expressed genes (DEGs) were identified using the Generalized linear model and ratio likelihood tests of the EdgeR package, according to its manual. Differentially expressed genes were defined as those genes with an absolute logFC (i.e. logarithmic fold change) > 1 and a Benjamini-Hochberg adjusted p-value as a significance threshold.

### 2.2 Patient clustering & analysis

Clusters of patients with similar expression profiles were identified using the expression data of the 185 psoriatic patient samples. The optimal number of clusters was decided based on the consensus between several widely-used metrics, namely the Elbow method, the Silhouette method and hierarchical clustering, and analyses were performed with the *fviz_nbclust* function from the “*factoextra*” package version 1.0.7 (Kassambara and Mundt, 2020). The clustering was done using the genes that among every psoriasis patient were varying the most. This was done to consider the dissimilarities between the psoriasis patients. The “*genefilter*” package version 1.74.1 (Gentleman et al., 2022) was utilized for finding these most variable genes.

To classify the psoriasis patients into clusters, the patient proximity matrices were calculated using Random Forest (RF) and Partitioning Around Medoids (PAM). These methods are available through the packages “*randomForest*” version 4.6.14 (Cutler and Wiener, 2022) and “*cluster*” version 2.1.2 (Maechler et al., 2022), respectively.

The 200 most variable genes across the patient cohort were considered for the clustering. Genes associated with the patients’ sex, and pseudogenes, were removed, resulting in a final list of 179 genes which was used for patient classification. In the PAM clustering algorithm, the psoriasis patient proximity matrix created by RF was used as an input with the specification of the number of clusters (k) to be identical to the optimal cluster number found earlier (k=2).

After the partition of the patients into two clusters, an investigation of potential differences between the clusters was performed. Sample metadata provided by Federico et al. were used to find potential patterns in the separation of patients. Additionally, the 179 most variable genes used to classify the patients were annotated for their reported role in psoriasis and association with disease severity. An additional differential expression analysis between the two patient clusters was performed to identify statistically significant differences in gene expression, using Cluster 1 as the baseline. A pathway enrichment analysis was performed to identify the processes separating the two clusters, using the ReactomePA package version 1.36.0 (Yu and He, 2016), with a false discovery rate (FDR) < 0.05.

### 2.3 Logical modeling

The PsoKC logical model published previously (Tsirvouli et al., 2021), representing psoriatic KCs, was used as the basis for building an extended model that represented the pathways that differentiate the two identified patient clusters. The initial model included the signaling pathways that are activated in psoriatic KCs upon activation of the main psoriatic cytokines (IL-17, IL-22, IFNγ and TNFα) and/or Prostaglandin E2 (PGE2) signaling. Psoriasis-specific behaviors of the KCs can be observed by incorporating “marker genes”: model entities involved in signaling that leads to specific phenotypes, such as abnormal apoptotic and proliferative patterns. Immune cell activation (for instance the activation of Th17 cells, neutrophils, or Th1 cells) as a result of cytokine and chemokine production by psoriatic KCs can also be deduced from marker gene specifications.

For the model extension, a selection of the pathways from the pathway enrichment analysis was gathered from SIGNOR 2.0 (SIGnaling Network Open Resource) (Perfetto et al., 2016) or from literature curation. To decide which of the genes from the enriched pathways were going to be included in the model, the pathway constituents were manually examined for having a reported role in psoriasis, and more specifically in the context of psoriatic KCs. The interactions from SIGNOR were also inspected for their relevance, and interactions solely from experimental data of human cells were considered. Model extensions were performed in GINsim version 3.0 (Gonzalez et al., 2006). During the construction of the model, the logical rules and regulatory interactions were iteratively refined until the model’s local and global predicted states corroborated the known biological observations of psoriatic KCs reported in the literature. An OR-operator was employed if the literature did not state otherwise. Each newly added node was annotated with its corresponding Uniprot ID and PubMed ID(s) that confirmed their function in psoriasis pathology and implication in KC biology.

A stable state analysis was performed using the bioLQM tool, which is part of the *Consortium for Logical Models and Tools* (CoLoMoTo) notebook (Naldi et al., 2018). The full analysis is available through a Jupyter notebook at https://github.com/Eirinits/psoKC2.0_model.

### 2.4 Model personalization

To explore possible differences in the development of psoriasis and response to perturbation among the two patient clusters, the PROFILE pipeline was employed (Béal et al., 2021). The PROFILE pipeline creates patient-specific profiles based on omics data obtained from these patients and then performs MaBoSS simulations to calculate the probability of a state (or phenotype) to be reached, among others. The gene expression data of the 185 patients were supplied to the pipeline, which transforms the data into continuous values between 0 and 1, by normalizing them. The normalization process depends on the distribution of the data, and is automatically defined and carried out by the tool. The normalized data are then used as rates of transition (activation/inactivation) for each node and for the probability of a node to be active or inactive at the start of the simulations (initial states).

Such transition rates and initial states were defined for each patient, by using the “*Soft Node Variant*” method described for PROFILE (Béal et al., 2019). The resulting 103 node activities and rates for each patient were then used to employ a set of MaBoSS simulations. To simulate intrinsic variation in KC signaling between patient clusters, the model’s inputs (i.e. cytokine and PGE2 receptors) were set to be active in the simulations. Additionally, to avoid “predisposing” patients to reach a certain phenotype, rates and probabilities for the marker nodes were excluded from the personalized patient profiles. To facilitate this analysis, the model was expanded with phenotype nodes, which can be ‘activated’ by the model’s marker genes. The simulations were performed in unperturbed models (wild type (WT) model) and in models where each non-marker node was considered a potential drug target. The latter simulations were aimed to identify potential differences between treatment responses.

## 3. Results

### 3.1 Patient cluster identification and characterization

For identifying clusters of patients based on their gene expression profiles, batch-corrected, normalized gene expression data from 185 patients and nine different studies were used. The 200 most variable genes among patients were selected for this classification. The complete annotation of these most variable genes with respect to their role in psoriasis or their link to severity can be found in Supplementary Table 2. Among these most variable genes, 21 genes were identified as linked to patient sex or were pseudogenes and therefore excluded from the analysis, to avoid the inclusion of known confounding factors. The remaining genes were subjected to an unsupervised classification using Random Forest (RF) dissimilarity matrices after which partitioning around medoid (PAM) clustering was performed, resulting in two patient clusters with 83 and 102 individuals, respectively.

These two clusters were next explored with respect to their correlation to the clinical data available for each patient. Metadata for each patient were collected from each individual study, focusing on the inclusion criteria of the patients, their sex and Psoriasis Area and Severity Index (PASI) score. The PASI score is a metric commonly used by clinicians to assess the severity of the disease. A PASI score less than 5 is associated with mild disease, a score between 5 and 10 with moderate disease, and a PASI score above 10 denotes a severe disease (CADTH Common Drug Review, 2018). PASI score information was available for 141 out of the 185 patients. As shown in Figure 1B, most patients (∼ 60%) in our study cohort displayed moderate psoriasis, but both patient clusters appeared to contain patients of all possible severities, with no particular pattern. However, many of the patients with no PASI score annotations were classified in Cluster 1.

**Figure 1.**
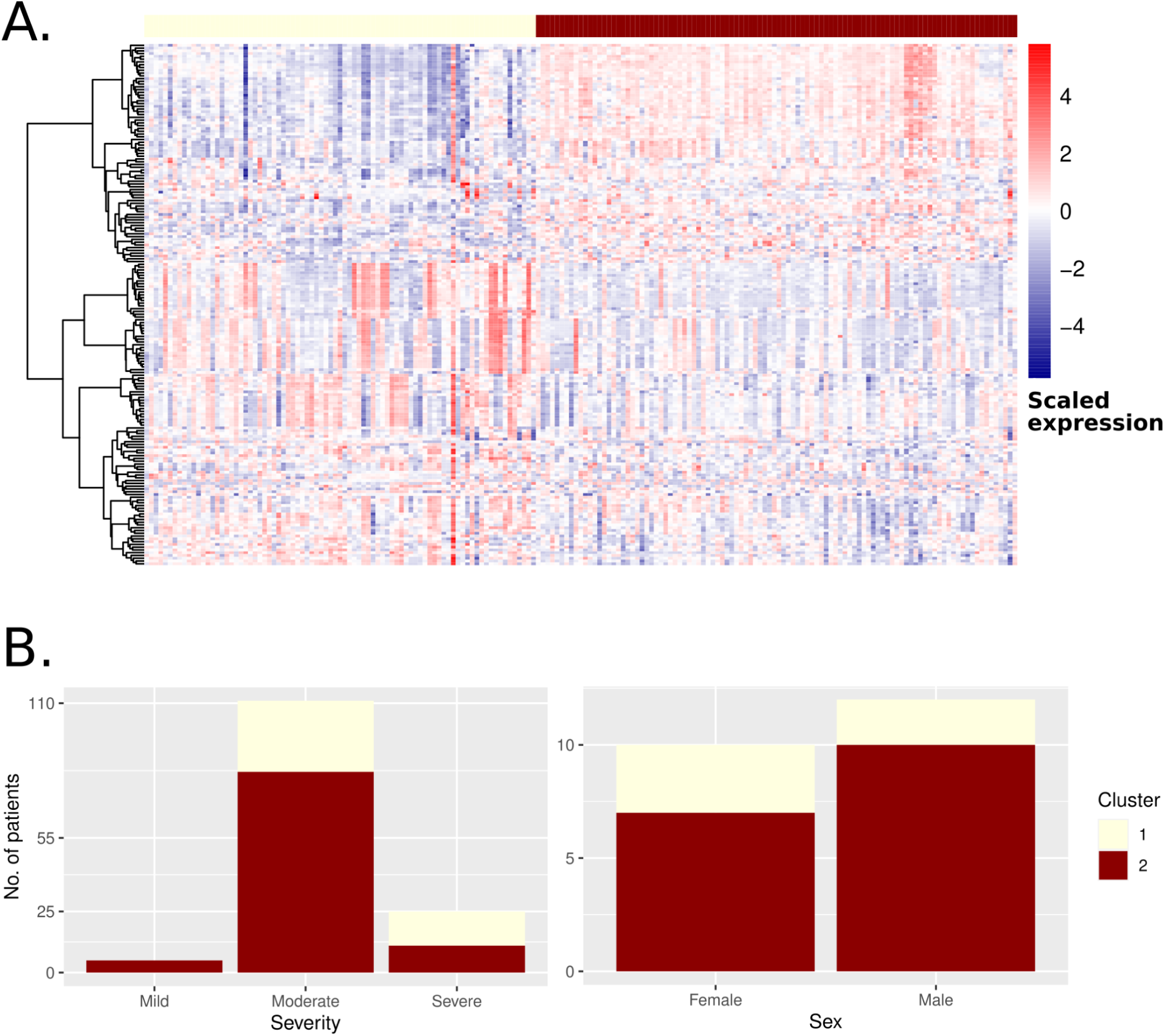
Characterization of the two identified patient clusters. (A) Heatmap of the scaled, normalized expression of the 185 most variable genes. Each column corresponds to a patient and each row to a gene. Patients are grouped according to their cluster classification, with light yellow representing Cluster 1 and dark red representing Cluster 2 (B) Stacked barplots representing the number of patients with different disease severity (left panel) and of a certain sex (right panel) in Cluster 1 (light yellow) and Cluster 2 (dark red).

Direct information about the patients’ sex was available only for 22 patients. Even within this limited sample, there was no clear association between sex and cluster, with both sexes assigned about equally to both clusters (Figure 1B). One of the included studies also specified whether the origin of the biopsy material was from the scalp, palmoplantar (i.e. palms and soles) or from the trunk and extremities. All but one of the scalp and palmoplantar samples were assigned to Cluster 1, while the rest of the samples from that study were separated over the two clusters. This might suggest that the two identified clusters could correspond to two different functional clusters of patients, as scalp and palmoplantar lesions were previously reported to be unresponsive to local and systemic therapies currently used to treat generalized psoriasis (Janagond et al., 2013).

While the two clusters exhibited differences in their expression profiles, they did not directly correlate with any of the available clinical variables. Some of the differences in expression levels between the two clusters concerned genes that have been linked to disease severity (Supplementary Figure 1), despite that both clusters contained patients of various severity levels. This highlights the difficulty of classifying patients into clinically meaningful subtypes using single omics data or without the identification of prognostic or severity biomarkers.

The poor association of the two gene expression clusters with the available clinical variables highlights the presence of alternative sources of variability in the patients’ expression, which could be associated with variable activity of biological pathways between the two clusters. To identify statistically significant differences in the expression between the two clusters, a differential expression analysis was performed and 117 genes were found significantly differentially expressed (FDR < 0.01). To further explore the biological processes that these DEGs are involved in, an enrichment analysis was performed separately on the upregulated and downregulated genes. Genes significantly upregulated in Cluster 2 were enriched for processes such as Interferon alpha/beta (IFNα/b) signaling, Interleukin-1 (IL-1) family signaling and Antimicrobial peptide (AMPs) signaling. On the other end, significantly downregulated genes were involved in processes related to Omega 3 and Omega 6 metabolism. Several studies have shown that decreased levels of circulating Omega 3 is correlated with disease severity and PASI score (Myśliwiec et al., 2017), although other studies remain inconclusive (Upala et al., 2017). Among the enriched processes, the term “Keratinization” was found both with over- and under-expressed genes. A closer look into the genes contributing to the enrichment of this term showed that proliferation markers of KCs, such as Keratin 6 (KRT6), were among the upregulated in Cluster 2 genes, while differentiation markers, such as Filagrin (FLG), were among the downregulated genes. Indeed, KCs overexpress KRT6 when in a highly activated and proliferative stage and under pathological conditions such as psoriasis (Zhang et al., 2019). FLG plays a significant role in the terminal differentiation of the epidermis and its role in the development of psoriasis has been related to its location in the psoriasis-susceptibility locus 4 (PSORS4), and its reduced expression in psoriatic skin has been described as a response to several of the main drivers of psoriasis, such as IL-17 (Zhang et al., 2017).

### 3.2 Model extension based on identified processes

To exploit the information contained in the two patient clusters, the processes and entities that distinguish the two clusters were used for constructing a logical model that would adequately represent the patients of the study. This was carried out by extending the psoriatic keratinocyte model (PsoKC model) that we published previously (Tsirvouli et al., 2021). Based on DEG and overrepresentation analysis, four main pathways were selected for model extension: IL-1 family signaling, with a focus on IL-1β and IL-36, and IFNα/β signaling. A fifth pathway, that of IL-6 signaling, was also included as it has a known role in disease severity and development, and it is involved in downstream regulation of several of the genes that separate the two patient clusters (e.g., Serpinb3 and Serpinb4 genes).

Several members of the IL-1 family are documented in literature to have mutations and variants that correlate with severe psoriasis. An example of such a gene is IL36RN, whose mutations and expression levels correlate with degree of acuteness in certain clinical variants of psoriasis (Sugiura, 2014). More generally, IL-36 members are involved in the amplification of general inflammation by regulating several cell types in the skin. Specifically in KCs, IL-36 promotes the release of inflammatory cytokines and chemokines and interferes with differentiation of the epidermis (Madonna et al., 2019). IL-1β is another member of the IL-1 family, which plays an orchestrating role in inflammatory responses. Dysregulation of IL-1β signaling is related to several chronic inflammatory diseases, including psoriasis (Cai et al., 2019). Certain IL-1β polymorphisms have been linked to the graveness of psoriasis, and its expression has been related to disease development and treatment responses (Cai et al., 2019).

IFNα/β signaling is the second signaling pathway that has been linked to more severe psoriasis and its dysregulation in patients was described as a susceptibility factor for the disease (Zhang, 2019). Type I interferons regulate several aspects of disease development, especially early initiation events by the activation of dendritic cells. In KCs, IFNα/β regulate the overexpression of IL-22 receptor genes, which leads to the increased responsiveness of KCs to the cytokine (Zhang, 2019, 1).

Lastly, the IL-6 pathway was included in the model extension due to its reported role in disease severity, mainly attributed to its direct effect on the activation of STAT3 and in consequence on KC hyperproliferation (Saggini et al., 2014). Recent studies have associated IL-6 levels to the risk of psoriatic arthritis development for psoriasis patients (Sobolev et al., 2022)

To keep the model to a manageable size and avoid the explosion of the state space, during the inclusion of the above pathways it was decided to focus on the entities whose expression changes between the two patient clusters and with regulatory connections to existing nodes of the base PsoKC model. For each of the pathways, a module containing the new nodes and their connection with the nodes of the PsoKC model is shown in Supplementary Figure 2.

### 3.3 The extended model - PsoKC 2.0

The PsoKC 2.0 model resulting from these extensions (Figure 2A.) consists of 135 nodes and 260 edges. The represented processes and nodes include pathways related to cell fate decision of KCs in response to their cytokine environment and pathways that can regulate disease severity either by inducing KC hyperproliferation or amplifying inflammatory responses. In order to assess the KC response to the different stimuli (or inputs), 66 nodes serve as markers that inform about the physiological state of KCs (e.g., whether a KC is proliferating or differentiating, or whether it secretes inflammatory cytokines to its environment that further promotes inflammation). Of these 66 nodes, 24 were added during the model extension. The annotation of each marker with its associated state can be found in Supplementary Table 3.

**Figure 2.**
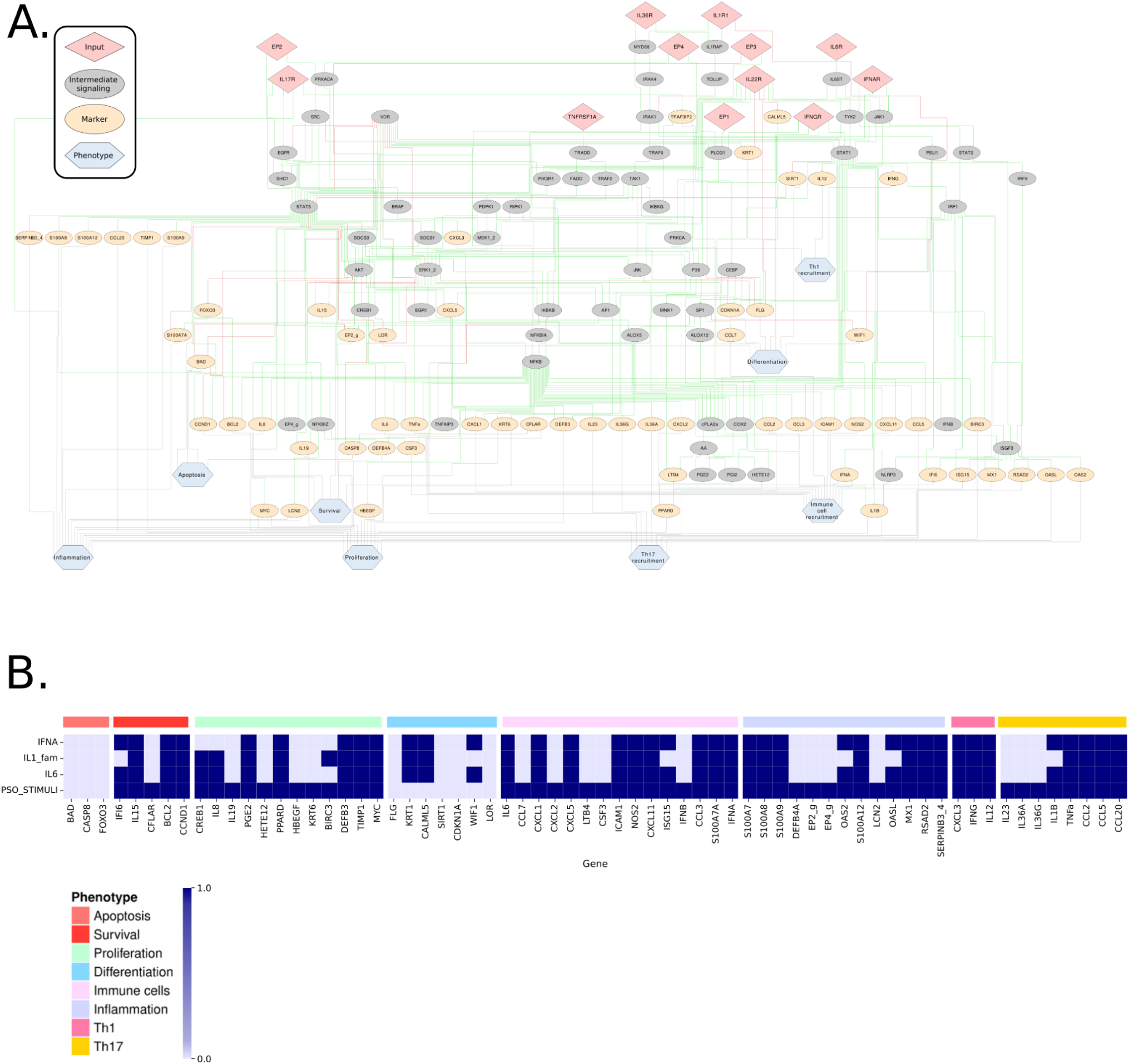
The PsoKC 2.0 model and its response to disease stimuli. A. The PsoKC 2.0 model representing psoriatic patient keratinocyte signaling, upon stimulation by psoriatic cytokines and prostaglandin E2 (PGE2). Each node represents a biological entity (i.e. proteins, fatty acids or genes), and each edge represents a regulatory interaction between two entities. Green arrows represent activating interactions and red arrows represent inhibiting interactions. Gray arrows show associations between markers and phenotypes. B. Heatmap of the stable states reached by the PsoKC 2.0 model. Each row represents a condition and each column a marker. Columns are organized based on the phenotypes they are promoting. The marker’s activity is represented by colors. Dark blue represents active nodes in the respective stable state, and light blue represents inactive nodes.

### 3.4 Model validation

After adding the five pathways, the final model was validated based on its ability to reproduce known behavior of psoriatic KCs under different conditions. The first validation was based on the same psoriatic conditions that the original model was validated against, and a second validation included literature-based evidence of the general behavior of the system and of individual markers.

Initially, the response of KCs to the newly added pathways was simulated separately for each pathway (IFNA, IL1_fam, IL6 conditions for the presence of IFNα, IL-1b and IL-36 simulation, and IL-6, respectively). Then, a condition characteristic of the psoriatic stimuli that KCs respond to was simulated with all the cytokines above set to be present (PSO_STIMULI condition).

As shown by Tsirvouli et al., the EP2 and EP4 receptors are upregulated and active in psoriatic KCs, therefore, they were set accordingly in these PsoKC 2.0 simulations. To verify this with the newly added pathways, simulations with and without the activation of the EP2/4 receptors were performed. The results were then compared against the available literature. All markers reached their expected state only if EP2 and EP4 receptors were set to be active in the simulated conditions (Supplementary Table 4 and 5), with the exception of IL-23 and IL-36A/G in the IL1_fam simulation. Therefore, the two receptors were set to active in all the downstream analysis. The results of the stable state analysis are presented in Figure 2B.

To verify that the newly added pathways were incorporated in a way that they accurately represent psoriatic signaling, the activity of the downstream markers of each pathway was checked against available literature. This confirmed that the model predicted the correct state of all markers except one (i.e. IFNβ). The predicted states of the markers and their literature-derived activities are presented in Supplementary Tables 6-7. All literature-derived activities are annotated with their respective PubMed ID.

In the presence of all psoriatic stimuli (PSO_STIMULI condition), the model reached the expected phenotype: The KCs are hyperproliferative with all survival and proliferation markers active, differentiation is impaired, and they secrete in their microenvironment cytokines and chemokines that further amplify inflammatory responses and recruit immune cells.

A comparison of their predicted activities from Tsirvouli et al. and the current model was also performed (Supplementary Table 8). A few discrepancies were observed in specific markers when simulating the individual effect of Th1- and Th17-derived cytokines. For example, the states of IL-19, CSF3 and DEFB4A nodes were predicted as active in the extended model in the presence of Th1 cytokines, while the opposite state was predicted by the PsoKC model. These discrepancies can be traced back to the inclusion of more pathways and the update of the logical rules of main regulators of the model. In the case of IL-19, CSF3 and DEFB4A, these are all regulated by the NFKBIZ transcription factor. While in the PsoKC model the NFKBIZ node is activated solely by IL-17R, the extended model includes the activating regulation of NFKBIZ by STAT3 and NF-κB, in response to the stimulation of IL-1 family cytokines (Müller et al., 2018). This indicates that the additional pathways help to represent aspects of the disease that were not fully captured by the PsoKC model.

### 3.5 Model personalization

After the validation of the model’s ability to recapitulate psoriasis development, the model was ready to be used to predict differences in the responses of the two patient clusters. This was done with the PROFILE pipeline (Béal et al., 2019), which creates expression profiles of individual patients. The pipeline uses RNA-seq data to infer the activation and inactivation rates of each node. This inference runs under the assumption that the higher the expression of a gene, the higher the probability is of a corresponding node to be activated. The addition of such rates to a Boolean model allows the semi-quantitative simulation of disease development and response to treatment, which can be observed as probability density plots of the activation of the model’s phenotypes.

As the model contains several markers that inform about the state of KCs, new phenotypic nodes were added to the model to facilitate the interpretation of the results of this analysis. Phenotypic nodes, as the name implies, are nodes that represent the different phenotypes and physiological states of KCs that can be represented in the model (Proliferation, Apoptosis, and Inflammation, but also Th1, Th17 and other immune cell recruitment). Phenotypic nodes are connected to the model in a way that they are activated by markers that are most representative or directly related to that phenotype. For example, both IL-19 and CCND1 are considered markers for proliferation. IL-19 promotes KC proliferation by stimulating fibroblasts to produce KGF. The CCND1 node, however, represents cyclin D, which is directly involved in proliferation. Therefore, only CCND1 was considered in the logical rules of the phenotypic nodes. For most of the phenotypes, the marker genes that promote the phenotype are linked with an “OR” boolean operator. For the differentiation phenotype, the markers that are related to differentiation are linked with an “AND” operator, as all markers are expected to be activated if a KC is terminally differentiated.

Two main analyses were performed with the personalized models: an analysis exploring the activation levels of each phenotypic node between the two clusters, and a perturbation analysis. The perturbation analysis aimed to identify perturbations that could lead to a differential treatment response between the patient clusters.

#### 3.5.1 Disease development simulations

To simulate the differences in the phenotype activation between the two clusters, an initial state was chosen where all psoriatic stimuli (*see* PSO_STIMULI condition in stable state analysis) were set to be active. For each subgroup, the results of the simulations are presented as density plots, with the node’s activity score on the x-axis (ranging from 0 to 1) and the proportion of patients with the respective node’s activity on the y-axis.

As seen in Figure 3, the probabilities of each phenotype to be active or inactive vary between the two clusters, with the figure panels showing a diverse dysregulation of phenotypic states. Apoptosis is predicted to be very significantly reduced in both groups, whereas the probability of proliferation seems to be somewhat higher in Cluster 2. Differences in the activation probabilities of inflammation- and immune-response related phenotypes are also shown. Interestingly, inflammatory markers and neutrophil-recruiting agents have a markedly higher probability of being active in Cluster 1, whereas markers related to the recruitment of immune cells, such as Th1 and Th17 cells, have a much higher probability of being activated in Cluster 2. The markers that are activating the “Inflammation” phenotype include entities as antimicrobial peptides and S100 peptides. These markers are mostly associated with the activation of innate immunity and general inflammatory responses (Hawkes et al., 2017). The balance between innate and adaptive immunity responses was previously proposed as a factor that influences the phenotypic characteristics of psoriasis (Schön, 2019). Following the onset of the disease, patients with a higher activation of innate immunity have a highly inflammatory psoriasis and flaring, while patients with a higher level of adaptive immunity exhibit a more stable, chronic disease (Schön, 2019). It is important to note that while the current model seems to capture some of these disease related phenotypic differences, it still represents the behavior of an individual cell type. Detailed intercellular aspects would be more accurately represented by multiscale models that are able to represent intercellular interactions.

**Figure 3.**
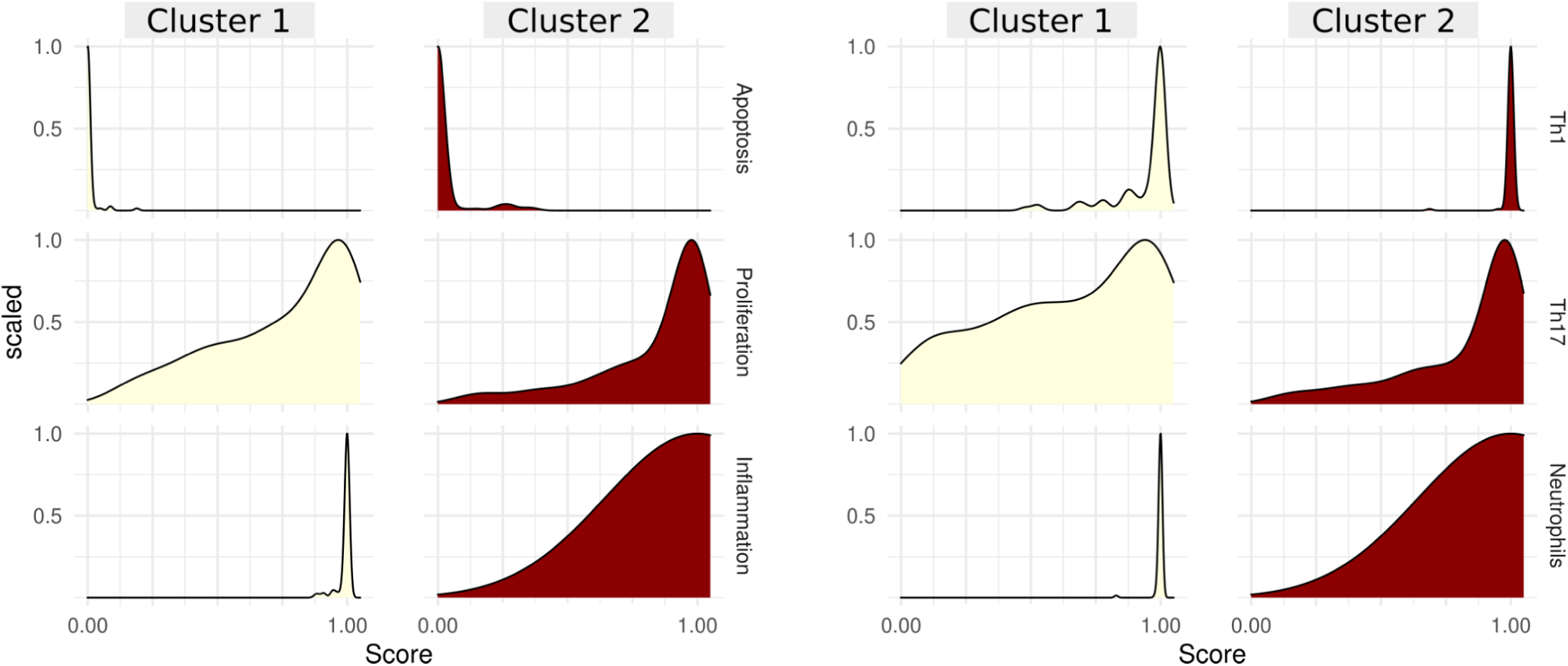
Density plots of the scores of keratinocyte cell fate phenotypes and immune response phenotypes that exhibit differences between the two patient clusters. The x-axis represents the probability that a phenotypic node is activated. The y-axis represents the scaled density of the distribution of the probability of activation across patients.

Based on the above, it is suggested that Cluster 1 could represent patients with overall higher levels of inflammation or inflammatory flares, while Cluster 2 shows a more strong promotion of adaptive immunity cell recruitment and survival, which is related with chronic psoriasis.

Since the newly added pathways have a role in psoriasis severity, the ability of the model to represent differences between severity groups was also assessed. This analysis was limited to the 141 patients with available PASI information. When patients are grouped together based on their PASI scores, the model captures the expected differences in the manifestation of the disease as shown in Figure 4. As expected, the probability of apoptosis and terminal differentiation are relatively higher in the mild subgroup than in the moderate and severe subgroups. Similarly, the probability of proliferation is high but still reduced compared to the moderate and severe subgroups. For the inflammation and immune phenotypes, the mild group shows a more variable response with some patients exhibiting reduced probabilities of activated inflammatory and immunostimulatory phenotypes. Patients with severe psoriasis are also predicted to have distinct phenotypes with fully inhibited apoptosis and differentiation, and a much higher probability of activation of all the other phenotypes. Patients with moderate disease show an intermediate profile, showcasing that the PsoKC 2.0 model is able to efficiently capture the variation in the activation of the pathways it contains.

**Figure 4.**
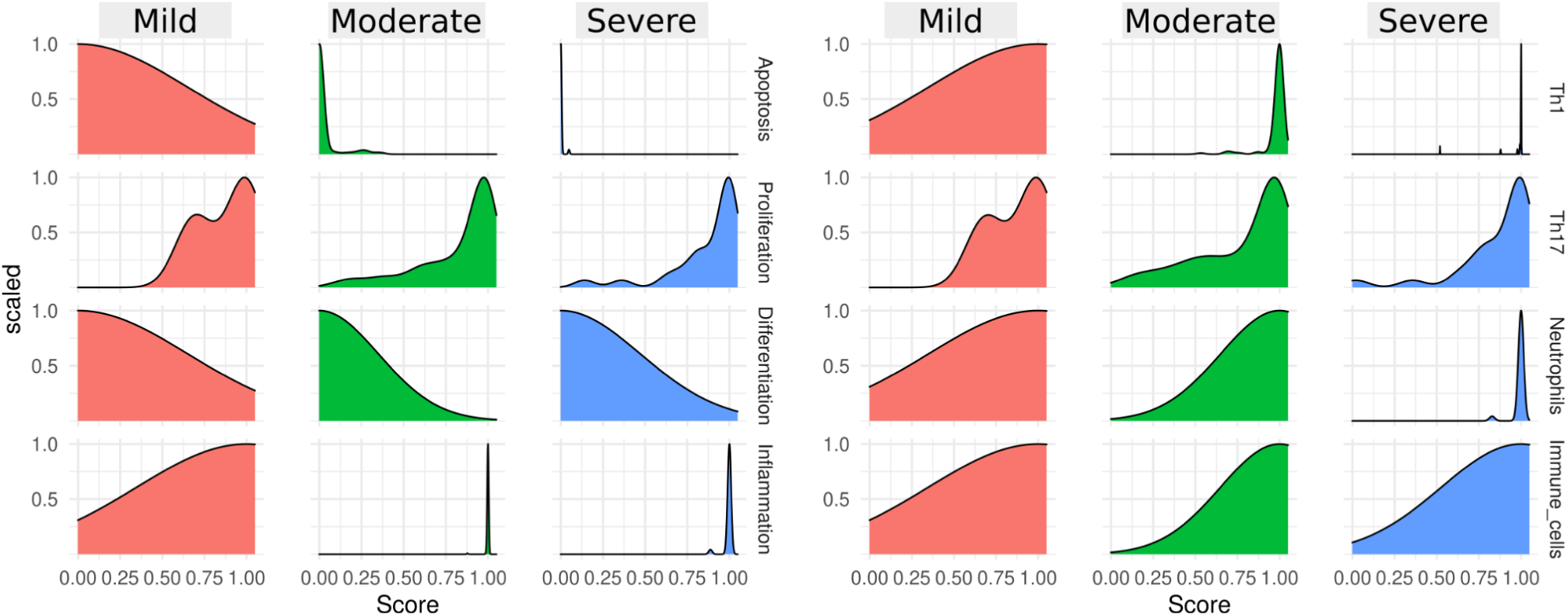
Density plots of the scores of keratinocyte cell fate phenotypes and immune response phenotypes in the 141 patients with available PASI score. The x-axis represents the probability that a phenotypic node is activated. The y-axis represents the scaled density of the distribution of the probability of activation across patients.

#### 3.5.2 Treatment responses

To explore the potential differences in response to treatment across patients, a perturbation analysis was performed in which the possible effect of biologics that aim to mitigate the effects of specific cytokines was simulated. In these simulations the model’s psoriatic stimuli were blocked. The *in silico* treatment outcome was quantified based on the PROFILE-predicted probabilities of the different phenotypes against the probabilities in untreated (WT) conditions as control. Based on the differences between the treated versus WT conditions, three levels of responses were defined: no responses: where the difference in probabilities is less than 0.1; low responses: where the difference in probabilities is less than 0.5; and strong responses, where the difference in probabilities exceeds 0.5. The density plots of all perturbation responses can be found in Supplementary Figures 3-14.

In general, responses were highly variable among patients, even within patients of the same gene expression cluster or PASI severity group. The most variable responses were observed among the apoptotic, proliferation, and Th cell promoting phenotypes. Three perturbations stood out as the most effective ones: the inhibition of IL-17, the inhibition of TNFα and the inhibition of PGE2 signaling. All three perturbations resulted in the reduction of proliferation and the Th-17 promoting phenotype.

The inhibition of IL-17 led to the reduction of proliferation in 179 of the 185 patient models. The extent of the reduction varied from 10% to 100%, with no particular pattern related to the two clusters or severity groups. The patients that showed a very low or no response to IL-17 inhibition were characterized by a relatively higher expression of several genes that include PLA2G4A, CREB1, JNK, MAPK3, BRAF, EGFR and IL6ST. JNK is associated with KC proliferation and the inhibition of KC differentiation (Gazel et al., 2006). IL6ST is a signaling transducer protein which was added to the model as part of the IL-6 pathway. IL6ST is activated downstream of IL-6 ligand-receptor binding and activates several inflammation-inducing and proliferation-promoting pathways (Piipponen et al., 2020). However, the expression of other survival promoting genes such as AKT1, NF-κB subunits, and MEK1, was lower in the non-/low-responders relative to the responding patients. Additionally, the same low or no-responding patients showed lower expression of TNFα receptor, TRADD, IRAK1, and TYK2. TYK2 is a key player in psoriasis and involved in the regulation of several inflammatory cascades in many of the cell types of the disease. In KCs, TYK2 regulates the activation of IkB-z, a regulator of NF-κΒ and STAT3 responses. Paradoxically, TYK2 appears to be expressed at lower levels in non-responders.

As the effect of the expression levels of individual genes is not so straightforward to interpret, additional simulations using MaBoSS were performed for one representative patient of each responder group (i.e. no response, low response, full response). The simulations were targeted to assess the trajectory of the activities of marker nodes and their regulators. There, the probability of NF-κB to be active, and by extension the probability of activation of the hyperproliferation marker KRT6, were inversely correlated with the strength of response. The full responder was the only patient where the activation of NF-κB was fully inhibited, whereas for the two other patients the probability of NF-κB and KRT6 was relatively low (∼ 0.1) but still not zero. Additionally, low responders exhibited a reduced probability of proliferation even in untreated settings, compared to the patients that showed a full response.

The inhibition of TNFα showed a high efficacy for reducing KC proliferation and the production of Th17-inducing ligands. However, there was no clear distinction between the expression profiles of responders and non-responders, as observed for the IL-17 inhibition. One of the genes that showed differences in expression between responders and non-responders to IL-17 inhibition was PLA2G4A, the gene coding for Cytosolic phospholipase A2α (cPLA2α). This phospholipase catalyzes the production of arachidonic acid (AA) and lysophosphatidylcholine (LPC). AA can be further metabolized to produce several fatty acids, such as PGE2. The inhibition of PGE2 signaling (in simulations encoded as the inactivation of the EP2 and EP4 receptors) resulted in the reduction of proliferation in 155 of the 185 patient models. Similar to the blocking of TNFα, the responders to the PGE2 inhibition did not separate from the non-responders, based on their expression patterns. The inhibition of individual EP receptors was not sufficient to reduce psoriatic phenotypes. This could be attributed to the shared downstream activation of signaling pathways, such as cAMP signaling (Rundhaug et al., 2011).

As one of the main problems with current treatment options for psoriasis is the lack of response of a small proportion of patients, the last analysis was targeted to those patients who were unresponsive or had limited responses to IL-17 inhibition but strongly responded to either PGE2 or TNFα, or both. These perturbation simulations showed that 10 patients with low or no response to IL-17 inhibition were responsive to PGE2 and/or TNFα blockade. In the whole cohort of patients these 10 showed the highest expression of genes such as CREB1, BRAF, SIRT1, EGFR and the lowest expression of AKT1, TYK2, IRAK1, and TRADD. The activity score assigned by PROFILE for the nodes in those patients where PGE2 and/or TNFα targeting could serve as an alternative to IL-17 inhibition is shown in Figure 5B.

**Figure 5.**
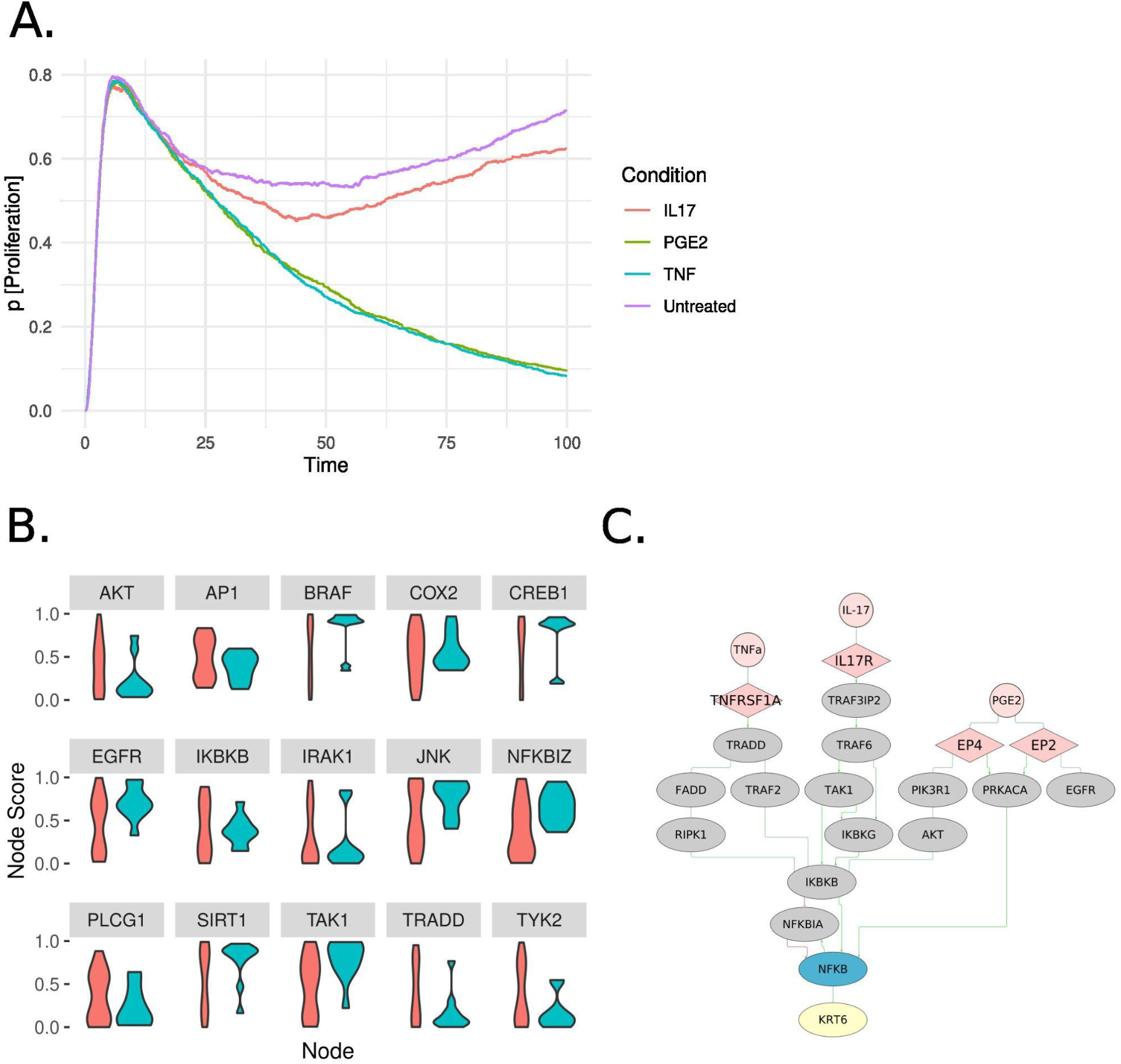
Treatment responses to the inhibition of IL-17, TNFα, and PGE2 differ among patients. A. Probability of an active proliferation (y-axis) in function of time (x-axis) in a patient that did not respond to IL-17 inhibition but responded to either the inhibition of PGE2 or TNFα. B. Violin plots of the model’s node scores for IL-17 low or non-responders, as generated by the PROFILE pipeline. Pink violins represent patients that did not respond to IL-17 but responded strongly to PGE2 and TNFα inhibition. Blue violins represent patients that did not show strong responses to any of the IL-17, PGE2 or TNFα inhibitions. Node scores correspond to the normalized expression of the gene that each node represents. A node with a higher score was set to have a higher probability of being active and a higher rate of activation in the model simulations. C. Schematic representation of NF-κB regulation downstream of the activation of IL-17, TNFα and PGE2 (i.e. EP2 and EP4) receptors, as represented in the PsoKC 2.0 model. Green arrows represent activating interactions and red arrows represent inhibiting interactions.

The patient ‘GSM1561614’ was used as an example to further analyze the responses to the different perturbations. This patient has moderate psoriasis and was classified as a ‘non-responder’ to IL-17 inhibition (reduction of proliferation < 0.01) and as a strong responder to both PGE2 and TNF inhibition. While the expression profile of this patient was not particularly distinct from the rest of the patients, it was characterized by a very high expression of NF-κB and its regulators. This high expression was translated into high probability of NF-κB initially active (∼90% probability) and getting activated along the course of the simulations.

To study in more detail the effects of perturbations, the state trajectories of individual nodes were compared against unperturbed simulations. The analysis was focused on intermediate signaling nodes of the model and the markers of KC cell fate. Many of the signaling nodes displayed a similar trajectory between all conditions, including the unperturbed one. Few nodes, such as NFKBIA for TNF inhibition, and CREB1 for PGE2 inhibition, differentiated their trajectory in only one condition.

Among the proliferation markers, KRT6 was the marker that followed a different trajectory between the perturbed conditions, with IL-17 inhibition resulting in a higher probability of activation of the hyperproliferation marker. In the model, KRT6 is regulated by two transcription factors, NF-κB and AP1. AP1 was not inhibited by any of the perturbations, therefore the differences in the activation of KRT6 could only be attributed to NF-κB. Indeed, IL-17 failed to fully inhibit NF-κB as opposed to PGE2 and TNFα inhibition. The regulation of the NF-κB as represented in the PsoKC 2.0 is shown in Figure 5C. Interestingly, PGE2 and TNFα contribute to the inhibition of NF-κB by two different mechanisms; With PGE2 signaling inhibited, the EP2 and EP4 receptors did not get activated and the downstream activation of cAMP-dependent protein kinase (PRKACA), and later of NF-κB, was inhibited. TNFα inhibition acted through the inhibition of RIPK1 and IKK-complex (represented in the model by one of its subunits, IKBKB). On the other hand, IL-17 did not (fully) inhibit any of the direct regulators of NF-κB. Combined with the high activity of NF-κB in the studied patient, IL-17 inhibition alone was unable to efficiently reduce KRT6 production and proliferation. It is important to note that this analysis cannot by itself support the causation of high NF-κB expression and negative responses to IL-17 inhibition. However, it provides an approximation of cases where the high activity of NF-κB and/or its regulators, caused by mutations or other dysregulations, could require alternative or combinatorial therapies to alleviate disease symptoms.

For the other perturbations, only very limited responses were predicted by the model for all the tested phenotypes. The lack of efficacy of those perturbations has been demonstrated also by *in vitro* studies or in clinical trials. More specifically, anti-IL-6 therapies have shown little or no effect in the treatment of the disease, and can be accompanied by several side effects, including the induction of psoriasis-like disease (Blauvelt, 2017). Similarly, IL-1β targeting was found to be ineffective in the treatment of chronic plaque-psoriasis (Tsai and Tsai, 2017). IL-22 has also been tested as a target but the current options failed to induce disease clearance (Tsai and Tsai, 2017). Lastly, the targeted cytokines act by stimulating KCs to produce a wide array of cytokines and chemokines that attract and promote the expansion of several immune cell types. Therefore, it could be expected that the effects of such perturbations, even if limited or moderate, would be reflected in the immune cell populations and not in KCs that the model represents.

## 4. Discussion

Although patients have seen an unprecedented improvement of treatment options against psoriasis, the disease remains incurable, and several patients fail to see complete clearance and suffer from side effects of varying severity. While several susceptibility genes have been identified as risk factor for developing psoriasis (Capon et al., 2007; Liu et al., 2008; Zhang et al., 2009) and for disease progression (Ramessur et al., 2022), effective biomarkers to classify patients into molecular subgroups or predict treatment outcome are scarce. A recently published review of systemic treatment response biomarkers reported several biomarkers, mostly predictive of the effect of TNFα agonists, ustekinumab (Anti-IL-12/23) and the immune system-suppressor methotrexate (Corbett et al., 2022). As the authors also denote, there is a lack of “hypothesis-free” biomarker testing, as most of the studies are focused only on molecular entities with a known role in psoriasis. Furthermore, well-defined molecular subtypes of plaque psoriasis with a predictive power on prognosis or treatment outcome are currently lacking. A few studies have tried to identify such subgroups using transcriptomics (Ainali et al., 2012; Krishnan and Kõks, 2022) or metabolomics (Dai et al., 2022) data. In our study, large scale transcriptomics datasets presented by Federico et al., were used to separate patients into clusters with similar gene expression profiles. The analysis proposed two main patient clusters. Each cluster had a distinct expression profile but, maybe surprisingly, this was not associated with any disease severity, as originally expected.

Our knowledge about dysregulated molecular mechanisms underlying diseases is becoming so extensive that the construction of computer models representing the disease not only becomes possible, but that it can also generate a useful tool for a detailed analysis of disease-related regulatory mechanisms and the discovery of possible treatment regimes. Previously we have published the PsoKC logical model that replicates the behavior of KCs dysregulated in psoriasis, and demonstrated a remarkable level of congruency between *in silico* perturbations and wet lab experiments performed on the same systems. Following the identification of two patient clusters, we have taken up the challenge to represent in a logical model in more detail the pathways and main regulatory components that seem to differ between the patient clusters. Looking at the genes displaying the largest differences among the patients, five pathways describing IFNα, IFNβ, IL-6, IL-1β and IL-36 signaling were added to the PsoKC model of psoriatic keratinocytes. The extended model (PsoKC 2.0) was validated by testing its ability to reproduce known biological observations of the disease in response to different stimuli.

The model was then used to identify potential functional differences between the two clusters by comparing the disease development and treatment responses between the two clusters. The model personalization pipeline PROFILE was used to create patient-specific expression profiles. There, nodes whose genes had higher expression were assigned to have a higher probability of being active and obtained a higher activation rate. This was assumed to approximate cases where patients have varying activity levels of certain molecular entities, causing distinct downstream effects for each patient or patient cluster. Simulations of the behavior of KCs in response to psoriatic stimuli showed that the personalized models were able to capture differential outcomes in disease progression between the two clusters. Patients of the first cluster showed a higher activation of innate immunity phenotypes, with a higher secretion of immunomodulatory ligands, such as antimicrobial peptides, while patients of the second cluster had a higher activation of adaptive immunity phenotypes, including the secretion of Th1 and Th17-recruiting and promoting cytokines and chemokines. The balance between innate and adaptive immunity can lead to different manifestations of the disease, where innate-dominant responses have a more highly inflammatory phenotype and adaptive-dominant responses a more stable and chronic phenotype (Schön, 2019). While additional validation would be needed to verify that the patients indeed displayed those differences in affected tissues, these observations highlight the value of model simulations for predicting how certain differences between patients, in our case differences in gene expression that on their own seem to binarize the patient cohort irrespective of disease severity, can be propagated in the system and lead to different outcomes. The processing of the gene expression data through logical modeling allows the application of regulary systems information to uncover and predict emergent phenotypes separating the two patient clusters and link such phenotypes to previously described manifestations of the disease.

Furthermore, as during the extension of the model many pathways with a documented role in disease severity were added, the model’s ability to separate between the development of mild, moderate and severe disease was evaluated. The grouping of the patients according to their PASI score confirmed that the model, coupled with omics data, can accurately capture the differences between severity groups and translate them into the expected differences in their physiological states.

Lastly, simulations targeted towards identifying differential treatment responses between patients or patient clusters were performed. Among the tested perturbations, three perturbations stood out as the most effective in reducing KC proliferation: IL-17 inhibition, TNFα inhibition and PGE2 inhibition. In all three perturbations, almost all patients were predicted to be responsive, with a reduced proliferation and secretion of Th17-related ligands. Patients with a predicted low or no response were not associated with any particular cluster or severity group.

The role of IL-17 and TNFα in several aspects of psoriasis and KC behavior has been described several times (Brembilla et al., 2018; Mylonas and Conrad, 2018). In KCs, IL-17 stimulation can promote their hyperproliferation and the release of cytokines, such as CCL20, that will ensure the survival, expansion and recruitment of immune cells (Furue et al., 2020). The importance of IL-17 in psoriasis was confirmed after the high efficacy of IL-17 inhibitors, such as secukinumab, ixekizumab, and brodalumab, in the treatment of moderate-to-severe cases (Silfvast-Kaiser et al., 2019). TNFα plays a key role in initiating events of psoriasis and acts as an amplifier of psoriatic responses by acting together with IL-17 and IL-22 in a synergistic manner (Chiricozzi et al., 2011; Grine et al., 2015). TNFα blockade was among the first biologics used for psoriasis and other inflammatory diseases, with a high efficacy and long term treatment of more severe cases of psoriasis (Schadler et al., 2019). *In vitro,* TNFα has been reported to regulate KC cell fate by regulating cell cycle and apoptotic genes (Banno et al., 2004). Several anti-IL-17 and anti-TNFα agents have shown promising results and have been approved for plaque psoriasis. IL-17 signaling can be inhibited at several levels, either with IL-17 antagonists (e.g Secukinumab), IL-17 receptor-A antagonists (e.g., Brodalumab) or via the double inhibition of IL-17A and IL-17F with Bimekizumab (Tsai and Tsai, 2017). For TNFα, four TNFα inhibitors have been approved for psoriasis, acting either by neutralizing soluble and/or membrane-bound TNFα (e.g Secukinumab and Infliximab) or by preventing interaction with its receptor (e.g., Adalimumab). In contrast, direct or indirect inhibitors of PGE2 have not been approved for psoriasis. PGE2 has a multifaceted role in the development of the disease: influencing the differentiation and effector functions of dendritic cells and Th cells (Chizzolini and Brembilla, 2009; An et al., 2021), stimulating KCs to secrete chemoattractants, and by dysregulating the normal life cycle of KCs (Tsirvouli et al., 2021). Inhibition of upstream regulators of PGE2 production such as cPLA2a (Ashcroft et al., 2020, 2) and COX-2 (Nițescu et al., 2022) have been described as promising for anti-psoriasis topical treatment, but further clinical studies are needed to assess the potentials of PGE2 signaling as a drug target in psoriasis.

Given the chronic nature of the disease and the need for life-long treatment, it is not uncommon that patients experience relapses or decreased treatment efficacy with time. In a few cases, patients might not respond at all or experience severe side effects, demanding the discontinuation of the treatment. For that, patients frequently have to switch between treatments or undergo a so-called rescue (or combination) therapy. In the studied cohort, 10 patients with a limited or no response to IL-17 inhibition were found to respond strongly to either PGE2 or TNFα inhibition. Additional simulations revealed that PGE2 and TNFα inhibition could fully inhibit NF-κB activity and KRT6 activation by inhibiting direct regulators of NF-κB. Patients not responding to IL-17 showed a higher expression of NF-κB among other prosurvival genes, which was encoded in the model as a higher activity of those entities. High NF-κB activity in dendritic cells was found to be predictive of the TNFα inhibitor, adalimumab (Andres-Ejarque et al., 2021), but no studies have shown a relation to anti-IL-17 effectiveness. Whether NF-κB activity can serve as a marker will require additional validation. The datasets used in this study consisted of bulk RNA-seq, which produces averaged gene expression of all the cell types and populations that are present in the sample. Bulk RNA-seq can give valuable insights on differences between samples and conditions (e.g., diseases versus healthy tissues) and highly (dys)regulated processes and pathways. However, it fails to accurately capture cell type-specific properties and heterogeneity. Therefore, technologies that address those limitations, such as single cell RNA-seq, possibly coupled with multicellular models, could provide more accurate predictions of treatment response biomarkers. Additionally, the integration of other omics data, especially of omics types that are more informative of a protein’s activity like phosphoproteomics, could aid the capturing of more variation between patients and patients groups.

Despite its limitations, the current study showcases how the integration of omics data and logical modeling can be used to predict differences in disease development and treatment response between patients, identify functional characteristics of groups of patients with a similar omics profile, identify potentially effective treatment alternatives to non-responder patients and help prioritize their testing and validation in the lab.

## Acknowledgements

The authors would like to acknowledge Laurence Calzone for her support on the use of the PROFILE pipeline and for providing us with an updated version of the pipeline. We would also like to thank Mari Løset for contributing with her clinical expertise to the initial parts of the project.

## Author Contributions

Conceptualization: MK, ET

Methodology: EA, ET

Data analysis, data analysis and model extension: EA

Model refinement, analysis and simulations: ET

Writing - Original draft: ET, EA

Writing - Review & Editing: ET, EA, MK

## Data Availability Statement

The study did not generate any new datasets. The code used in this study is fully available at https://github.com/Eirinits/psoKC2.0_model

## References

Ainali, C., Valeyev, N., Perera, G., Williams, A., Gudjonsson, J. E., Ouzounis, C. A., et al. (2012). Transcriptome classification reveals molecular subtypes in psoriasis. BMC Genomics 13, 472. doi: 10.1186/1471-2164-13-472.

Albanesi, C. (2019). “Immunology of Psoriasis,” in Clinical Immunology (Elsevier), 871–878.e1. doi: 10.1016/B978-0-7020-6896-6.00064-8.

Albanesi, C., Madonna, S., Gisondi, P., and Girolomoni, G. (2018). The Interplay Between Keratinocytes and Immune Cells in the Pathogenesis of Psoriasis. Front. Immunol. 9. doi: 10.3389/fimmu.2018.01549.

An, Y., Yao, J., and Niu, X. (2021). The Signaling Pathway of PGE2 and Its Regulatory Role in T Cell Differentiation. Mediators Inflamm. 2021, e9087816. doi: 10.1155/2021/9087816.

Andres-Ejarque, R., Ale, H. B., Grys, K., Tosi, I., Solanky, S., Ainali, C., et al. (2021). Enhanced NF-κB signaling in type-2 dendritic cells at baseline predicts non-response to adalimumab in psoriasis. Nat. Commun. 12, 4741. doi: 10.1038/s41467-021-25066-9.

Armstrong, A. W., and Read, C. (2020). Pathophysiology, Clinical Presentation, and Treatment of Psoriasis: A Review. JAMA 323, 1945–1960. doi: 10.1001/jama.2020.4006.

Ashcroft, F. J., Mahammad, N., Midtun Flatekvål, H., J. Feuerherm, A., and Johansen, B. (2020). cPLA2α Enzyme Inhibition Attenuates Inflammation and Keratinocyte Proliferation. Biomolecules 10, 1402. doi: 10.3390/biom10101402.

Banno, T., Gazel, A., and Blumenberg, M. (2004). Effects of tumor necrosis factor-alpha (TNF alpha) in epidermal keratinocytes revealed using global transcriptional profiling. J. Biol. Chem. 279, 32633–32642. doi: 10.1074/jbc.M400642200.

Béal, J., Montagud, A., Traynard, P., Barillot, E., and Calzone, L. (2019). Personalization of Logical Models With Multi-Omics Data Allows Clinical Stratification of Patients. Front. Physiol. 9. doi: 10.3389/fphys.2018.01965.

Béal, J., Pantolini, L., Noël, V., Barillot, E., and Calzone, L. (2021). Personalized logical models to investigate cancer response to BRAF treatments in melanomas and colorectal cancers. PLOS Comput. Biol. 17, e1007900. doi: 10.1371/journal.pcbi.1007900.

Blauvelt, A. (2017). IL-6 Differs from TNF-α: Unpredicted Clinical Effects Caused by IL-6 Blockade in Psoriasis. J. Invest. Dermatol. 137, 541–542. doi: 10.1016/j.jid.2016.11.022.

Blighe, K., Brown, A.-L., Carey, V., Hooiveld, G., and Lun, A. (2022). PCAtools: PCAtools: Everything Principal Components Analysis. Bioconductor version: Release (3.16) doi: 10.18129/B9.bioc.PCAtools.

Brembilla, N. C., Senra, L., and Boehncke, W.-H. (2018). The IL-17 Family of Cytokines in Psoriasis: IL-17A and Beyond. Front. Immunol. 9. Available at: https://www.frontiersin.org/articles/10.3389/fimmu.2018.01682 [Accessed December 12, 2022].

CADTH Common Drug Review (2018). CADTH Canadian Drug Expert Committee Recommendation: Guselkumab (Tremfya — Janssen Inc.): Indication: For the treatment of adult patients with moderate-to-severe plaque psoriasis who are candidates for systemic therapy or phototherapy. Ottawa (ON): Canadian Agency for Drugs and Technologies in Health Available at: http://www.ncbi.nlm.nih.gov/books/NBK532985/ [Accessed December 14, 2022].

Cai, Y., Xue, F., Quan, C., Qu, M., Liu, N., Zhang, Y., et al. (2019). A Critical Role of the IL-1β-IL-1R Signaling Pathway in Skin Inflammation and Psoriasis Pathogenesis. J. Invest. Dermatol. 139, 146–156. doi: 10.1016/j.jid.2018.07.025.

Capon, F., Di Meglio, P., Szaub, J., Prescott, N. J., Dunster, C., Baumber, L., et al. (2007). Sequence variants in the genes for the interleukin-23 receptor (IL23R) and its ligand (IL12B) confer protection against psoriasis. Hum. Genet. 122, 201–206. doi: 10.1007/s00439-007-0397-0.

Catapano, M., Vergnano, M., Romano, M., Mahil, S. K., Choon, S.-E., Burden, A. D., et al. (2020). IL-36 Promotes Systemic IFN-I Responses in Severe Forms of Psoriasis. J. Invest. Dermatol. 140, 816–826.e3. doi: 10.1016/j.jid.2019.08.444.

Chiricozzi, A., Guttman-Yassky, E., Suárez-Fariñas, M., Nograles, K. E., Tian, S., Cardinale, I., et al. (2011). Integrative Responses to IL-17 and TNF-α in Human Keratinocytes Account for Key Inflammatory Pathogenic Circuits in Psoriasis. J. Invest. Dermatol. 131, 677–687. doi: 10.1038/jid.2010.340.

Chizzolini, C., and Brembilla, N. C. (2009). Prostaglandin E2: igniting the fire. Immunol. Cell Biol. 87, 510–511. doi: 10.1038/icb.2009.56.

Corbett, M., Ramessur, R., Marshall, D., Acencio, M. L., Ostaszewski, M., Barbosa, I. A., et al. (2022). Biomarkers of systemic treatment response in people with psoriasis: a scoping review. Br. J. Dermatol. 187, 494–506. doi: 10.1111/bjd.21677.

Cutler, F. original by L. B. and A., and Wiener, R. port by A. L. and M. (2022). randomForest: Breiman and Cutler’s Random Forests for Classification and Regression. Available at: https://CRAN.R-project.org/package=randomForest [Accessed December 12, 2022].

Dai, D., He, C., Wang, S., Wang, M., Guo, N., and Song, P. (2022). Toward Personalized Interventions for Psoriasis Vulgaris: Molecular Subtyping of Patients by Using a Metabolomics Approach. Front. Mol. Biosci. 9. Available at: https://www.frontiersin.org/articles/10.3389/fmolb.2022.945917 [Accessed December 12, 2022].

Federico, A., Hautanen, V., Christian, N., Kremer, A., Serra, A., and Greco, D. (2020). Manually curated and harmonised transcriptomics datasets of psoriasis and atopic dermatitis patients. Sci. Data 7, 343. doi: 10.1038/s41597-020-00696-8.

Furue, M., Furue, K., Tsuji, G., and Nakahara, T. (2020). Interleukin-17A and Keratinocytes in Psoriasis. Int. J. Mol. Sci. 21. doi: 10.3390/ijms21041275.

Gazel, A., Banno, T., Walsh, R., and Blumenberg, M. (2006). Inhibition of JNK Promotes Differentiation of Epidermal Keratinocytes *. J. Biol. Chem. 281, 20530–20541. doi: 10.1074/jbc.M602712200.

Gentleman, R., Carey, V. J., Huber, W., and Hahne, F. (2022). genefilter: genefilter: methods for filtering genes from high-throughput experiments. Bioconductor version: Release (3.16) doi: 10.18129/B9.bioc.genefilter.

Gonzalez, A. G., Naldi, A., Sánchez, L., Thieffry, D., and Chaouiya, C. (2006). GINsim: A software suite for the qualitative modelling, simulation and analysis of regulatory networks. Biosystems 84, 91–100. doi: 10.1016/j.biosystems.2005.10.003.

Grine, L., Dejager, L., Libert, C., and Vandenbroucke, R. E. (2015). An inflammatory triangle in psoriasis: TNF, type I IFNs and IL-17. Cytokine Growth Factor Rev. 26, 25–33. doi: 10.1016/j.cytogfr.2014.10.009.

Hawkes, J. E., Chan, T. C., and Krueger, J. G. (2017). Psoriasis pathogenesis and the development of novel targeted immune therapies. J. Allergy Clin. Immunol. 140, 645–653. doi: 10.1016/j.jaci.2017.07.004.

Janagond, A. B., Kanwar, A. J., and Handa, S. (2013). Efficacy and safety of systemic methotrexate vs. acitretin in psoriasis patients with significant palmoplantar involvement: a prospective, randomized study. J. Eur. Acad. Dermatol. Venereol. JEADV 27, e384–389. doi: 10.1111/jdv.12004.

Kassambara, A., and Mundt, F. (2020). factoextra: Extract and Visualize the Results of Multivariate Data Analyses. Available at: https://CRAN.R-project.org/package=factoextra [Accessed December 12, 2022].

Krishnan, V. S., and Kõks, S. (2022). Transcriptional Basis of Psoriasis from Large Scale Gene Expression Studies: The Importance of Moving towards a Precision Medicine Approach. Int. J. Mol. Sci. 23, 6130. doi: 10.3390/ijms23116130.

Liu, Y., Helms, C., Liao, W., Zaba, L. C., Duan, S., Gardner, J., et al. (2008). A Genome-Wide Association Study of Psoriasis and Psoriatic Arthritis Identifies New Disease Loci. PLOS Genet. 4, e1000041. doi: 10.1371/journal.pgen.1000041.

Madonna, S., Girolomoni, G., Dinarello, C. A., and Albanesi, C. (2019). The Significance of IL-36 Hyperactivation and IL-36R Targeting in Psoriasis. Int. J. Mol. Sci. 20, 3318. doi: 10.3390/ijms20133318.

Maechler, M., original), P. R. (Fortran, original), A. S. (S, original), M. H. (S, Hornik [trl, K., maintenance(1999-2000)), ctb] (port to R., et al. (2022). cluster: “Finding Groups in Data”: Cluster Analysis Extended Rousseeuw et al. Available at: https://CRAN.R-project.org/package=cluster [Accessed December 12, 2022].

Michalak-Stoma, A., Bartosińska, J., Kowal, M., Juszkiewicz-Borowiec, M., Gerkowicz, A., and Chodorowska, G. (2013). Serum levels of selected Th17 and Th22 cytokines in psoriatic patients. Dis. Markers 35, 625–631. doi: 10.1155/2013/856056.

Müller, A., Hennig, A., Lorscheid, S., Grondona, P., Schulze-Osthoff, K., Hailfinger, S., et al. (2018). IκBζ is a key transcriptional regulator of IL-36–driven psoriasis-related gene expression in keratinocytes. Proc. Natl. Acad. Sci. 115, 10088–10093. doi: 10.1073/pnas.1801377115.

Mylonas, A., and Conrad, C. (2018). Psoriasis: Classical vs. Paradoxical. The Yin-Yang of TNF and Type I Interferon. Front. Immunol. 9. Available at: https://www.frontiersin.org/articles/10.3389/fimmu.2018.02746 [Accessed November 14, 2022].

Myśliwiec, H., Baran, A., Harasim-Symbor, E., Myśliwiec, P., Milewska, A. J., Chabowski, A., et al. (2017). Serum fatty acid profile in psoriasis and its comorbidity. Arch. Dermatol. Res. 309, 371–380. doi: 10.1007/s00403-017-1748-x.

Naldi, A., Hernandez, C., Levy, N., Stoll, G., Monteiro, P. T., Chaouiya, C., et al. (2018). The CoLoMoTo Interactive Notebook: Accessible and Reproducible Computational Analyses for Qualitative Biological Networks. Front. Physiol. 9. doi: 10.3389/fphys.2018.00680.

Nițescu, D. A.-M., Păunescu, H., Ștefan, A. E., Coman, L., Georgescu, C. C., Stoian, A. C., et al. (2022). Anti-Psoriasis Effect of Diclofenac and Celecoxib Using the Tail Model for Psoriasis. Pharmaceutics 14, 885. doi: 10.3390/pharmaceutics14040885.

Perfetto, L., Briganti, L., Calderone, A., Cerquone Perpetuini, A., Iannuccelli, M., Langone, F., et al. (2016). SIGNOR: a database of causal relationships between biological entities. Nucleic Acids Res. 44, D548–554. doi: 10.1093/nar/gkv1048.

Piipponen, M., Li, D., and Landén, N. X. (2020). The Immune Functions of Keratinocytes in Skin Wound Healing. Int. J. Mol. Sci. 21, 8790. doi: 10.3390/ijms21228790.

R Core Team (2013). R: A Language and Environment for Statistical Computing. R Foundation for Statistical Computing, Vienna. doi: http://www.R-project.org/.

Ramessur, R., Corbett, M., Marshall, D., Acencio, M. L., Barbosa, I. A., Dand, N., et al. (2022). Biomarkers of disease progression in people with psoriasis: a scoping review. Br. J. Dermatol. doi: 10.1111/bjd.21627.

Ran, D., Cai, M., and Zhang, X. (2019). Genetics of psoriasis: a basis for precision medicine. *Precis*. Clin. Med. 2, 120–130. doi: 10.1093/pcmedi/pbz011.

Ritchie, M. E., Phipson, B., Wu, D., Hu, Y., Law, C. W., Shi, W., et al. (2015). limma powers differential expression analyses for RNA-sequencing and microarray studies. Nucleic Acids Res. 43, e47. doi: 10.1093/nar/gkv007.

Robinson, M. D., McCarthy, D. J., and Smyth, G. K. (2010). edgeR: a Bioconductor package for differential expression analysis of digital gene expression data. Bioinformatics 26, 139–140. doi: 10.1093/bioinformatics/btp616.

Rundhaug, J. E., Simper, M. S., Surh, I., and Fischer, S. M. (2011). The role of the EP receptors for prostaglandin E2 in skin and skin cancer. Cancer Metastasis Rev. 30, 465–480. doi: 10.1007/s10555-011-9317-9.

Saggini, A., Chimenti, S., and Chiricozzi, A. (2014). IL-6 as a Druggable Target in Psoriasis: Focus on Pustular Variants. J. Immunol. Res. 2014, e964069. doi: 10.1155/2014/964069.

Schadler, E. D., Ortel, B., and Mehlis, S. L. (2019). Biologics for the primary care physician: Review and treatment of psoriasis. Dis. Mon. 65, 51–90. doi: 10.1016/j.disamonth.2018.06.001.

Schön, M. P. (2019). Adaptive and Innate Immunity in Psoriasis and Other Inflammatory Disorders. Front. Immunol. 10. doi: 10.3389/fimmu.2019.01764.

Silfvast-Kaiser, A., Paek, S. Y., and Menter, A. (2019). Anti-IL17 therapies for psoriasis. Expert Opin. Biol. Ther. 19, 45–54. doi: 10.1080/14712598.2019.1555235.

Sobolev, V. V., Denisova, E. V., Chebysheva, S. N., Geppe, N. A., and Korsunskaya, I. M. (2022). IL-6 Gene Expression as a Marker of Pathological State in Psoriasis and Psoriatic Arthritis. Bull. Exp. Biol. Med. 173, 77–80. doi: 10.1007/s10517-022-05497-0.

Sugiura, K. (2014). The genetic background of generalized pustular psoriasis: IL36RN mutations and CARD14 gain-of-function variants. J. Dermatol. Sci. 74, 187–192. doi: 10.1016/j.jdermsci.2014.02.006.

Tsai, Y.-C., and Tsai, T.-F. (2017). Anti-interleukin and interleukin therapies for psoriasis: current evidence and clinical usefulness. Ther. Adv. Musculoskelet. Dis. 9, 277–294. doi: 10.1177/1759720X17735756.

Tsirvouli, E., Ashcroft, F., Johansen, B., and Kuiper, M. (2021). Logical and experimental modeling of cytokine and eicosanoid signaling in psoriatic keratinocytes. iScience 24, 103451. doi: 10.1016/j.isci.2021.103451.

Upala, S., Yong, W. C., Theparee, T., and Sanguankeo, A. (2017). Effect of omega-3 fatty acids on disease severity in patients with psoriasis: A systematic review. Int. J. Rheum. Dis. 20, 442–450. doi: 10.1111/1756-185X.13051.

Yu, G., and He, Q.-Y. (2016). ReactomePA: an R/Bioconductor package for reactome pathway analysis and visualization. Mol. Biosyst. 12, 477–479. doi: 10.1039/C5MB00663E.

Zhang, L. (2019). Type1 Interferons Potential Initiating Factors Linking Skin Wounds With Psoriasis Pathogenesis. Front. Immunol. 10, 1440. doi: 10.3389/fimmu.2019.01440.

Zhang, X., Yin, M., and Zhang, L. (2019). Keratin 6, 16 and 17—Critical Barrier Alarmin Molecules in Skin Wounds and Psoriasis. Cells 8, 807. doi: 10.3390/cells8080807.

Zhang, X.-J., Huang, W., Yang, S., Sun, L.-D., Zhang, F.-Y., Zhu, Q.-X., et al. (2009). Psoriasis genome-wide association study identifies susceptibility variants within LCE gene cluster at 1q21. Nat. Genet. 41, 205–210. doi: 10.1038/ng.310.

Zhang, Y., Tu, C., Wang, S., and Xiao, S. (2017). Expression of Skin Barrier Protein Filaggrin in Skin Diseases without Atopic Dermatitis. J. Biosci. Med. 6, 101–112. doi: 10.4236/jbm.2018.61009.

