## SupplementaryMaterial for "Patient-specific logical models replicate phenotype responses to psoriatic and anti-psoriatic stimuli"

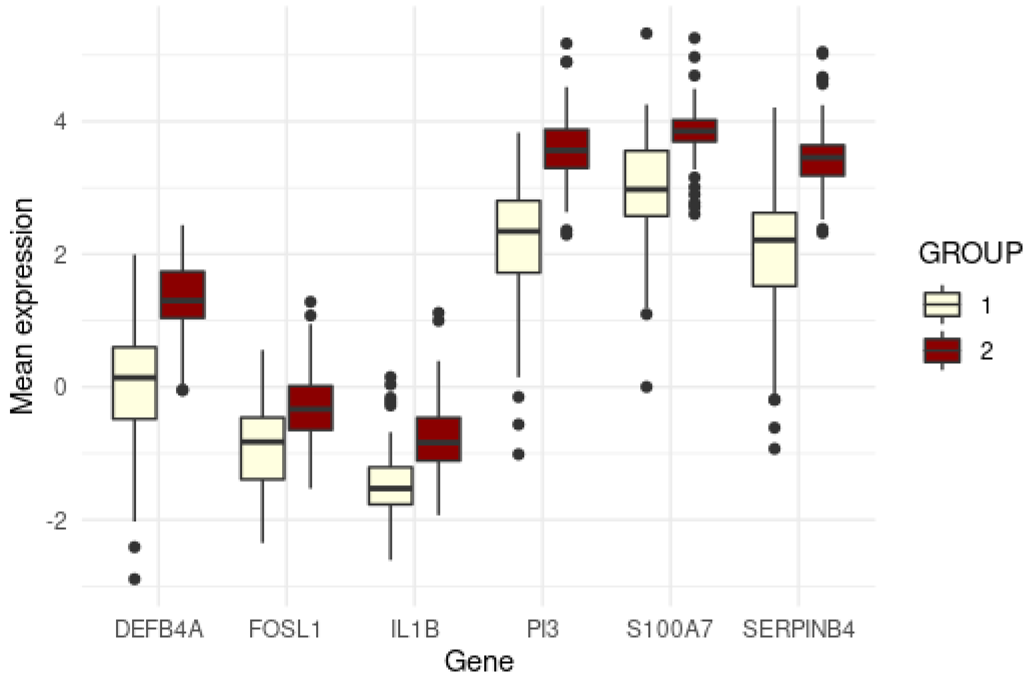

**Supplementary Figure 1.** Barplots of the scaled, mean expression of selected genes that positively correlate with psoriasis severity.

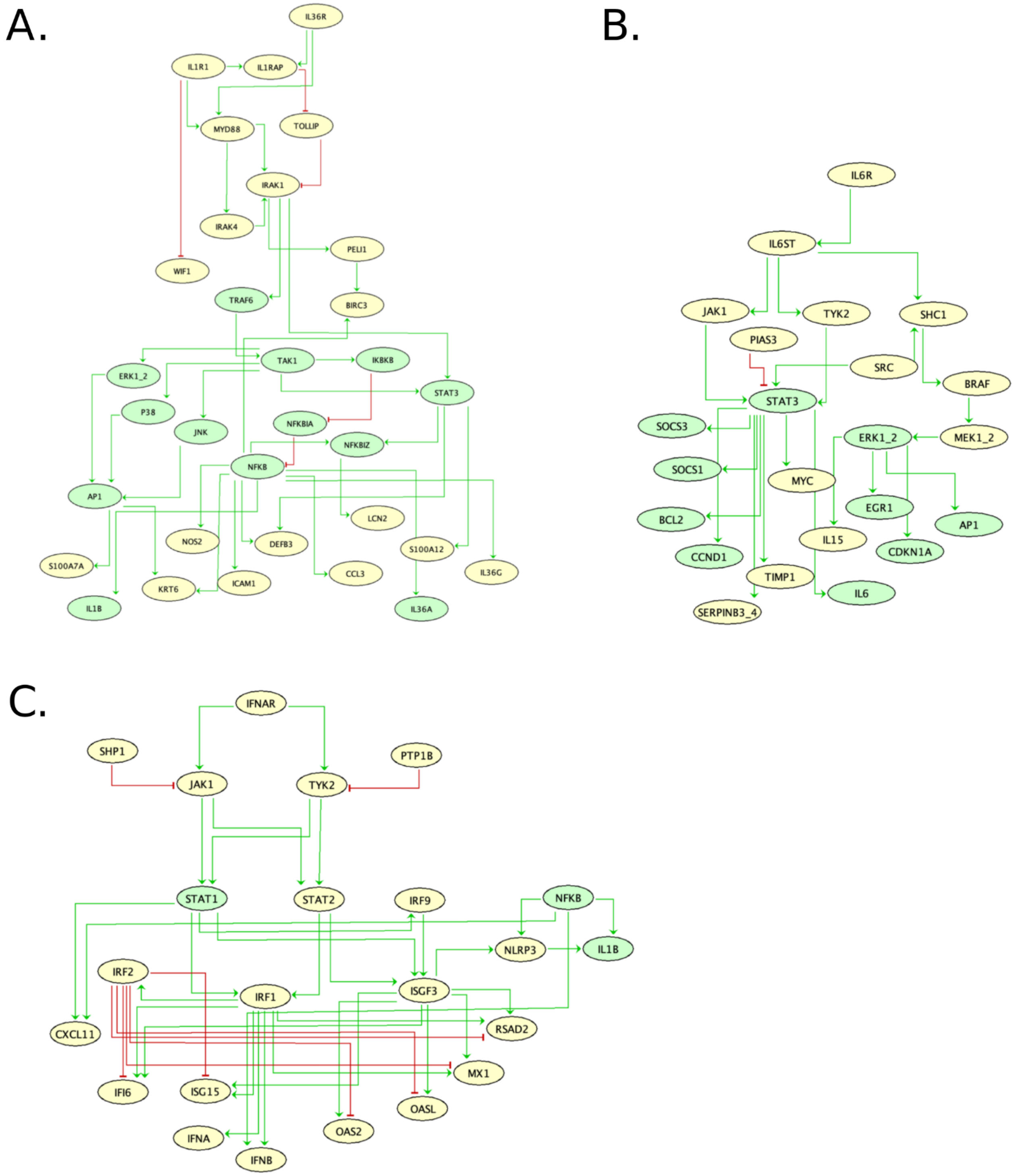

**Supplementary Figure 2.** A presentation of the pathways that were added in the base model (Tsirvouli et al., 2021). Yellow nodes present newly added nodes, while green nodes represent nodes that were already present in the base model. Green arrows represent activating interactions, and red arrows represent inhibiting interactions. A. IL-36 and IL-1β pathways. B. IL-6 pathway C. IFNα/β pathway

**Supplementary Figures 3-14.** Density plots of the scores of keratinocyte cell fate phenotypes and immune response phenotypes that exhibit differences between the two patient clusters. The x-axis represents the probability that a phenotypic node is activated. The y-axis represents the scaled density of the distribution of the probability of activation across patients.

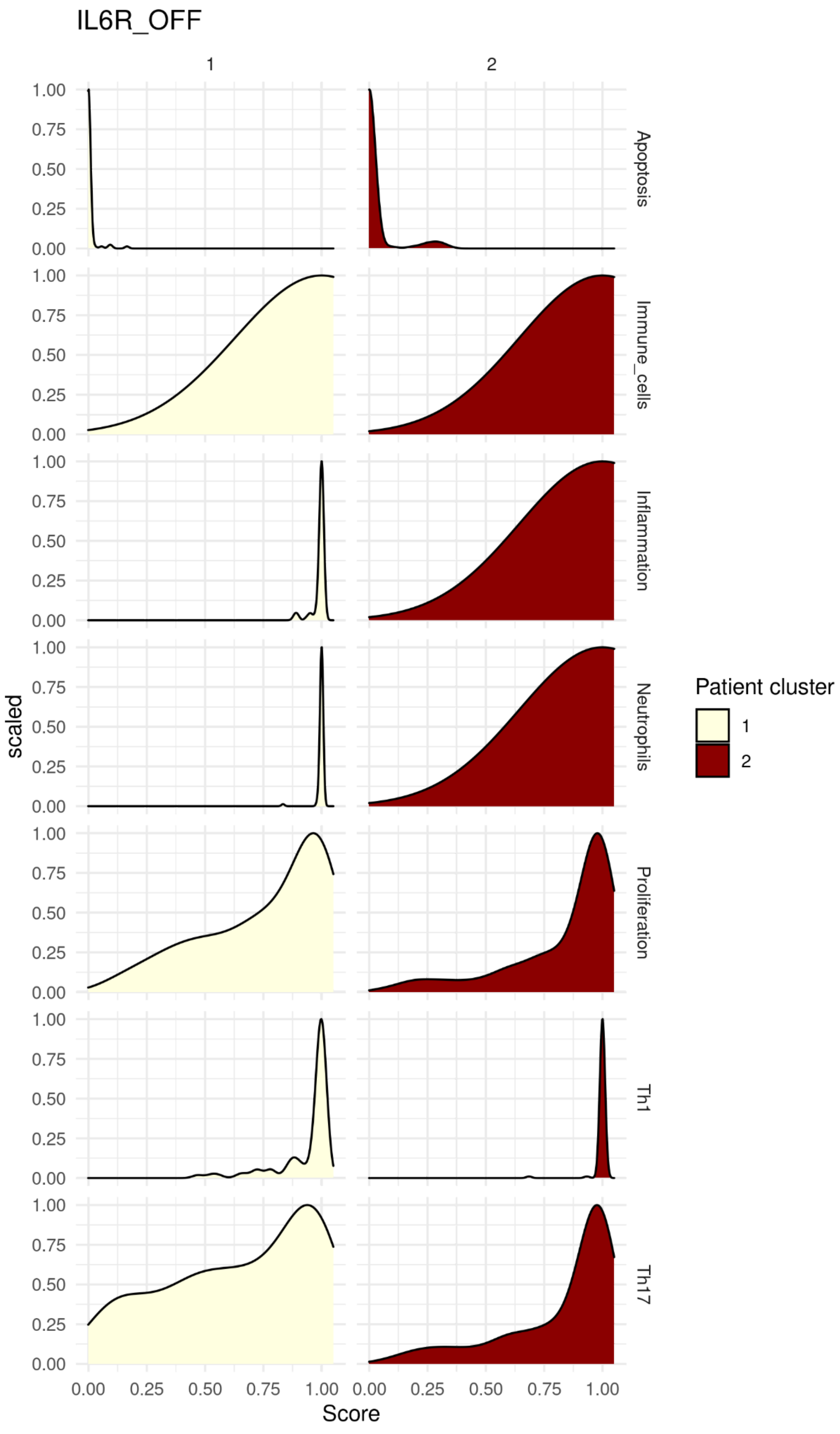

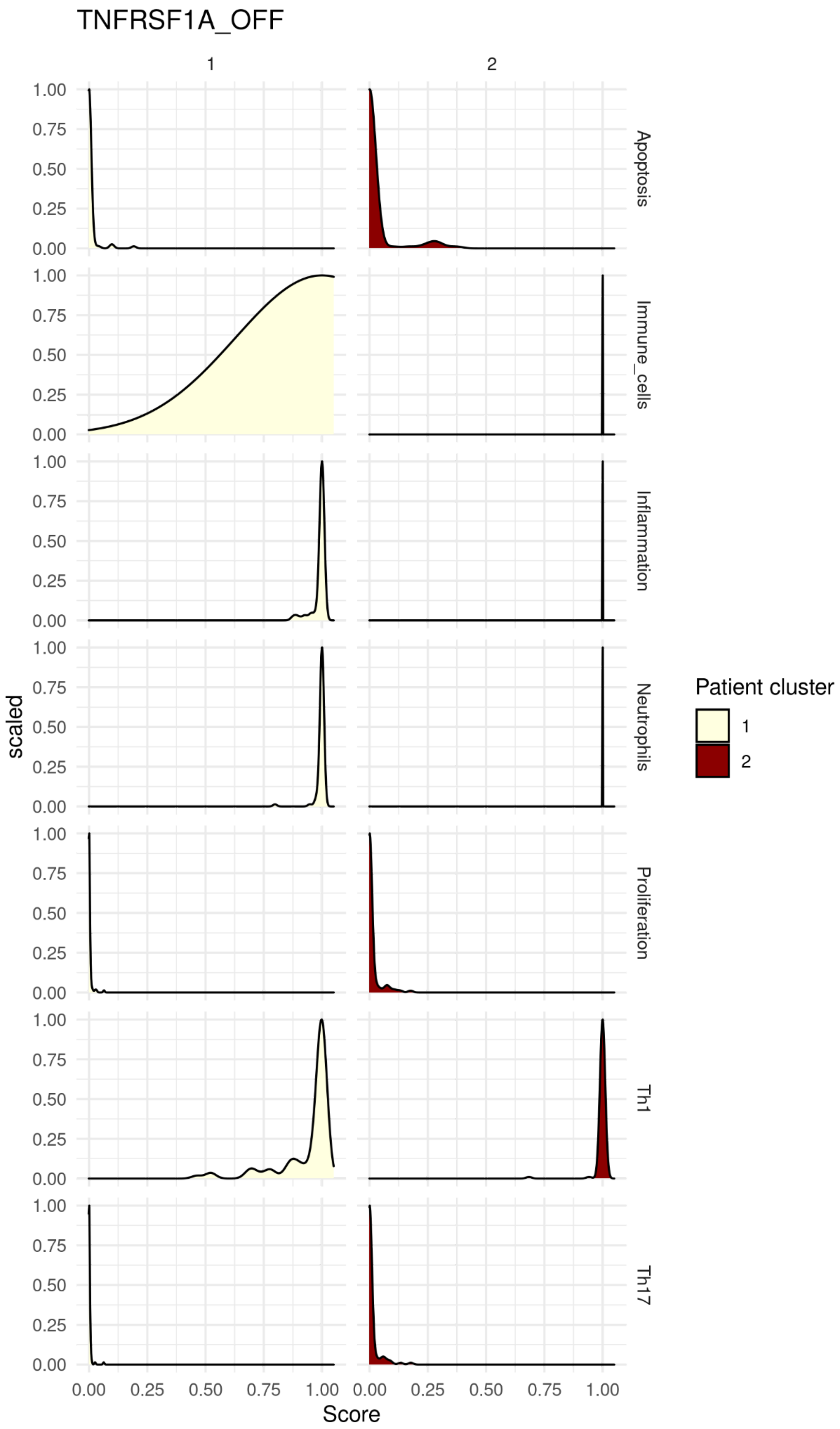

TNFRSF1A_OFF
1
2
Apoptosis
Immune_cells
Inflammation
Neutrophils
Proliferation
Th1
Th17
scaled
Score
0.00
0.25
0.50
0.75
1.00
Patient cluster
1
2

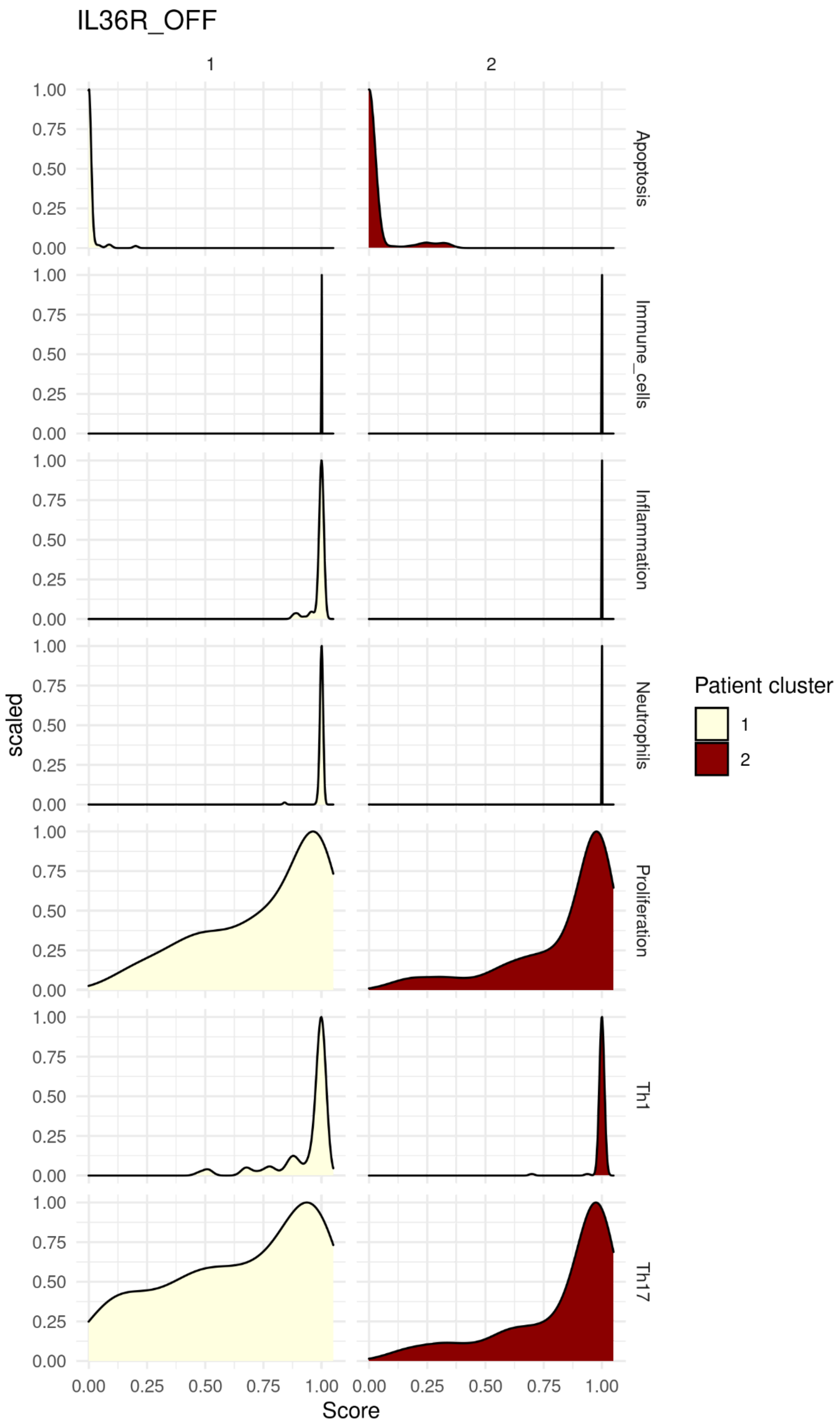
IL36R_OFF
1
2
Apoptosis
Immune_cells
Inflammation
Neutrophils
Proliferation
Th1
Th17
scaled
Score
0.00
0.25
0.50
0.75
1.00
Patient cluster
1
2

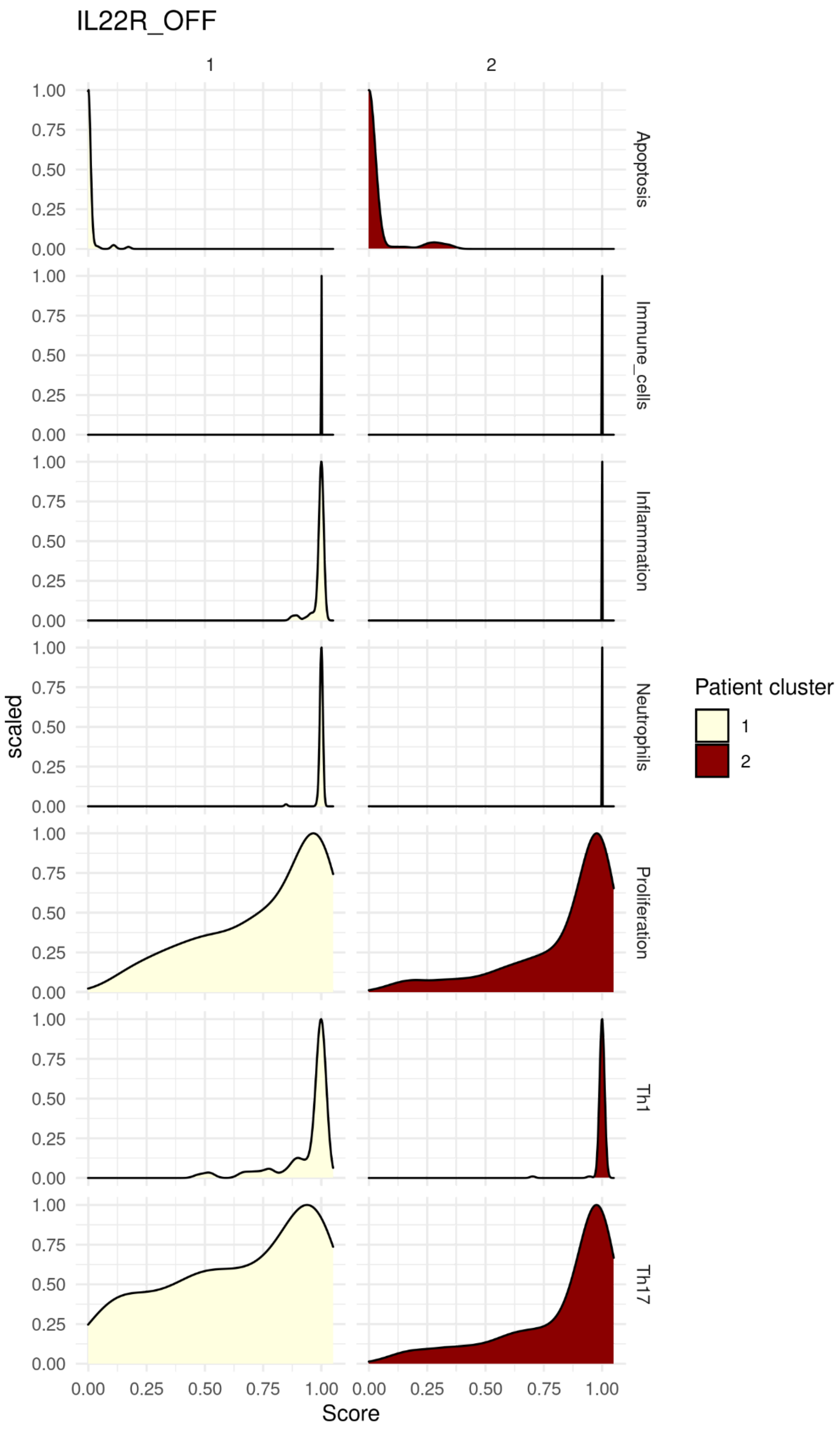
IL22R_OFF
1
2
Apoptosis
Immune_cells
Inflammation
Neutrophils
Proliferation
Th1
Th17
scaled
1.00
0.75
0.50
0.25
0.00
0.00 0.25 0.50 0.75 1.00
Score
Patient cluster
1
2

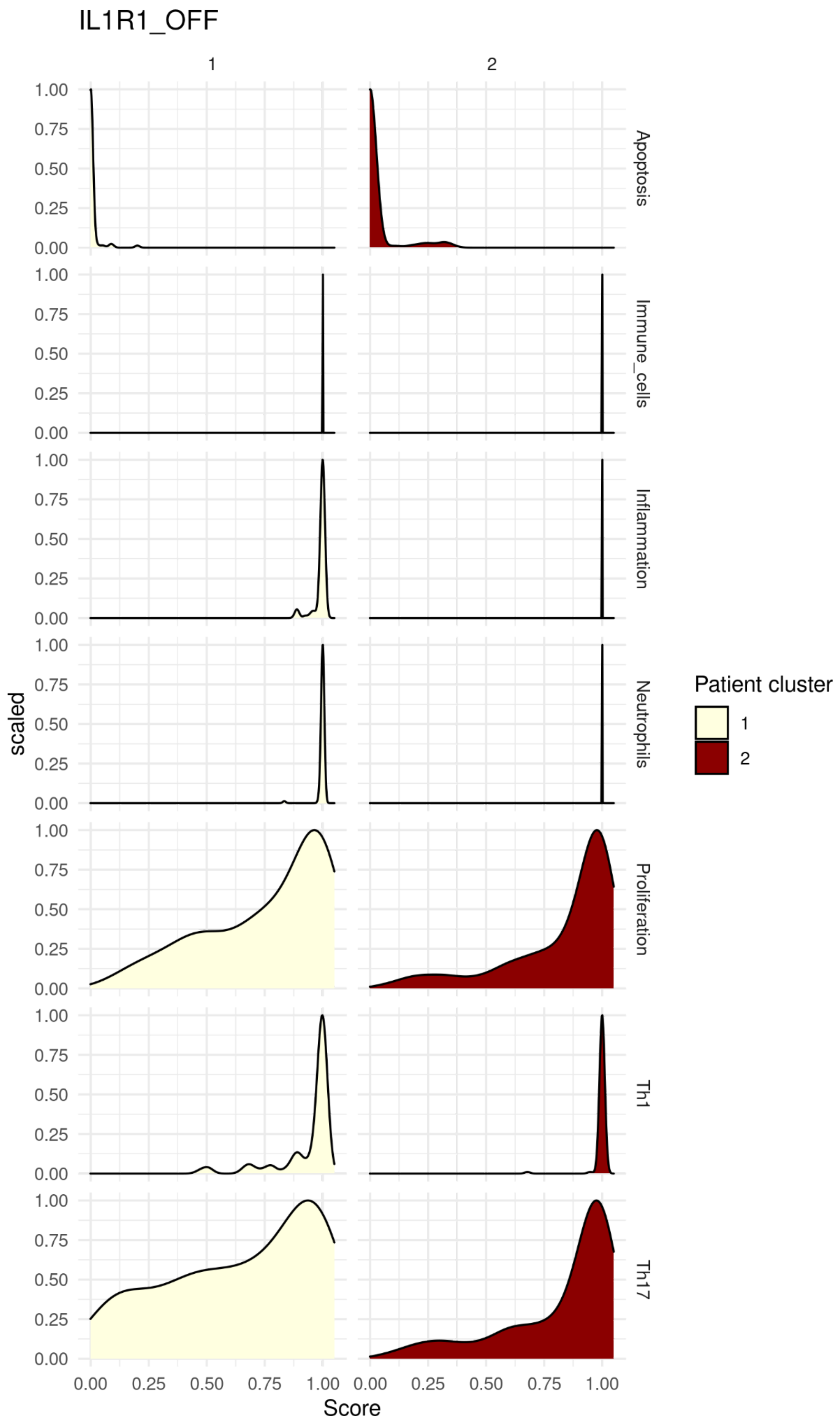

IL1R1_OFF
1
2
Apoptosis
Immune_cells
Inflammation
Neutrophils
Proliferation
Th1
Th17
scaled
Score
1.00
0.75
0.50
0.25
0.00
0.00
0.25
0.50
0.75
1.00
Patient cluster
1
2

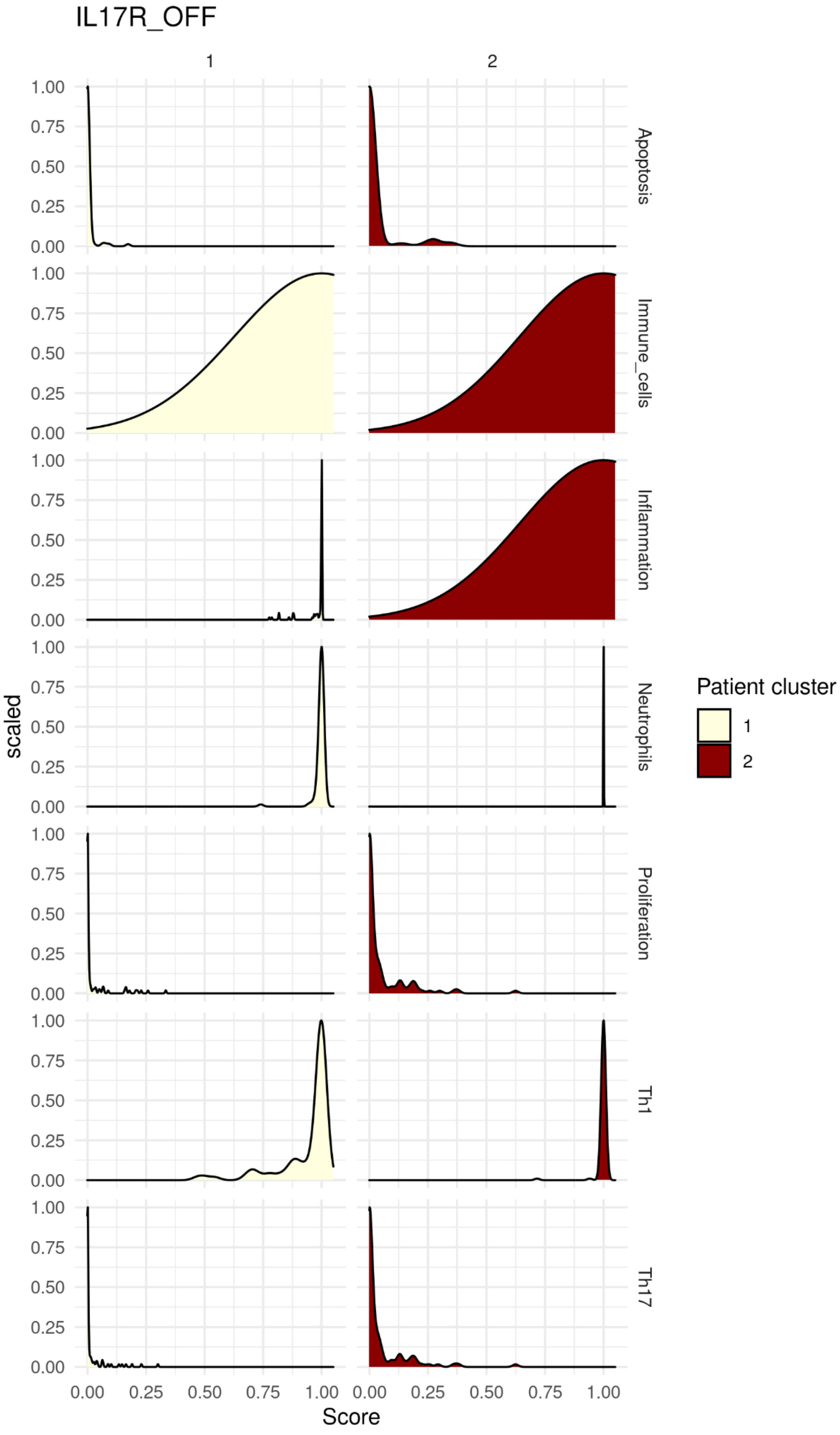

IL17R_OFF
1
2
Apoptosis
Immune_cells
Inflammation
Neutrophils
Proliferation
Th1
Th17
scaled
1.00
0.75
0.50
0.25
0.00
Score
0.00
0.25
0.50
0.75
1.00
Patient cluster
1
2

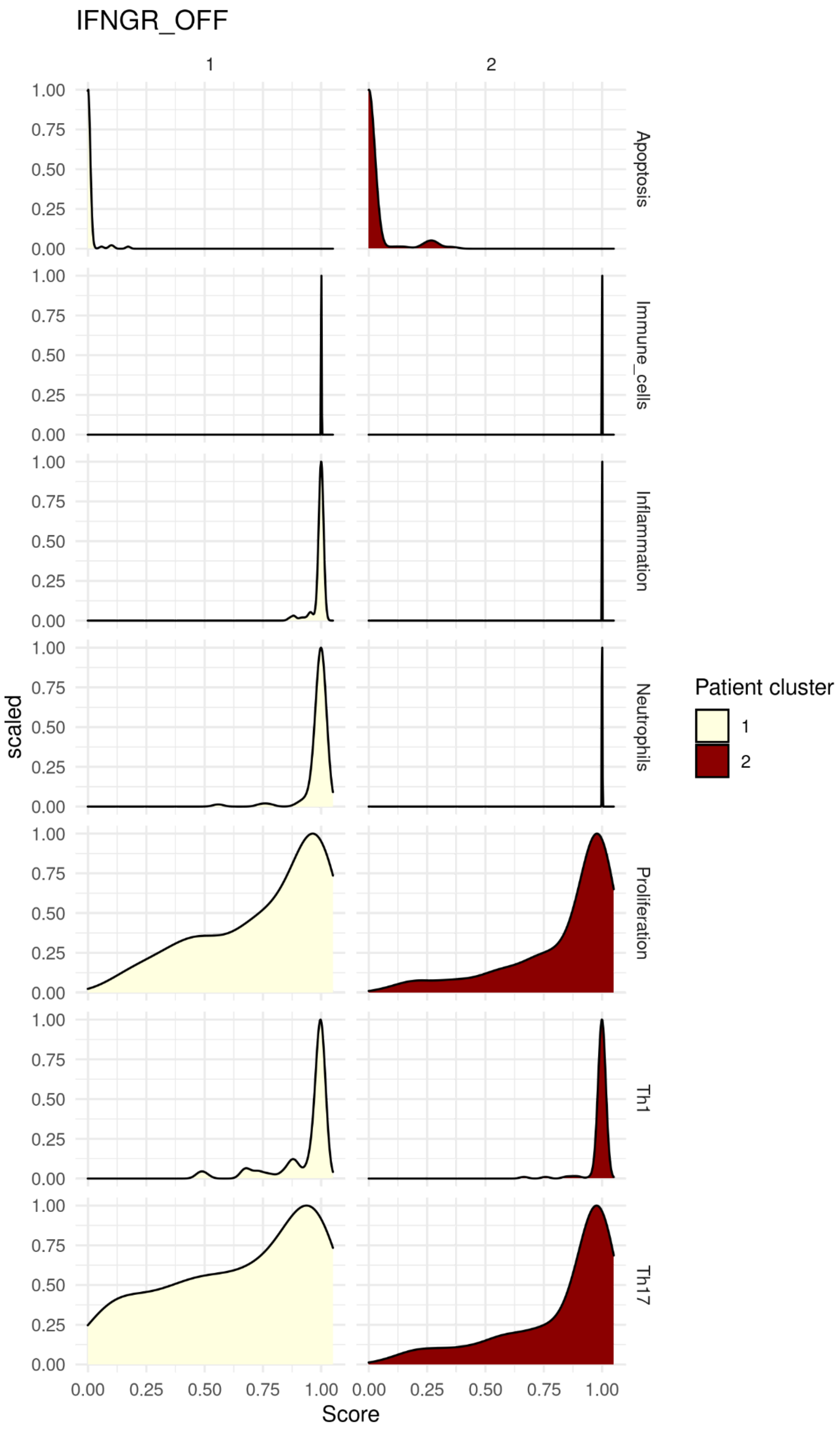

IFNGR_OFF
1
2
Apoptosis
Immune_cells
Inflammation
Neutrophils
Proliferation
Th1
Th17
scaled
Score
0.00
0.25
0.50
0.75
1.00
Patient cluster
1
2

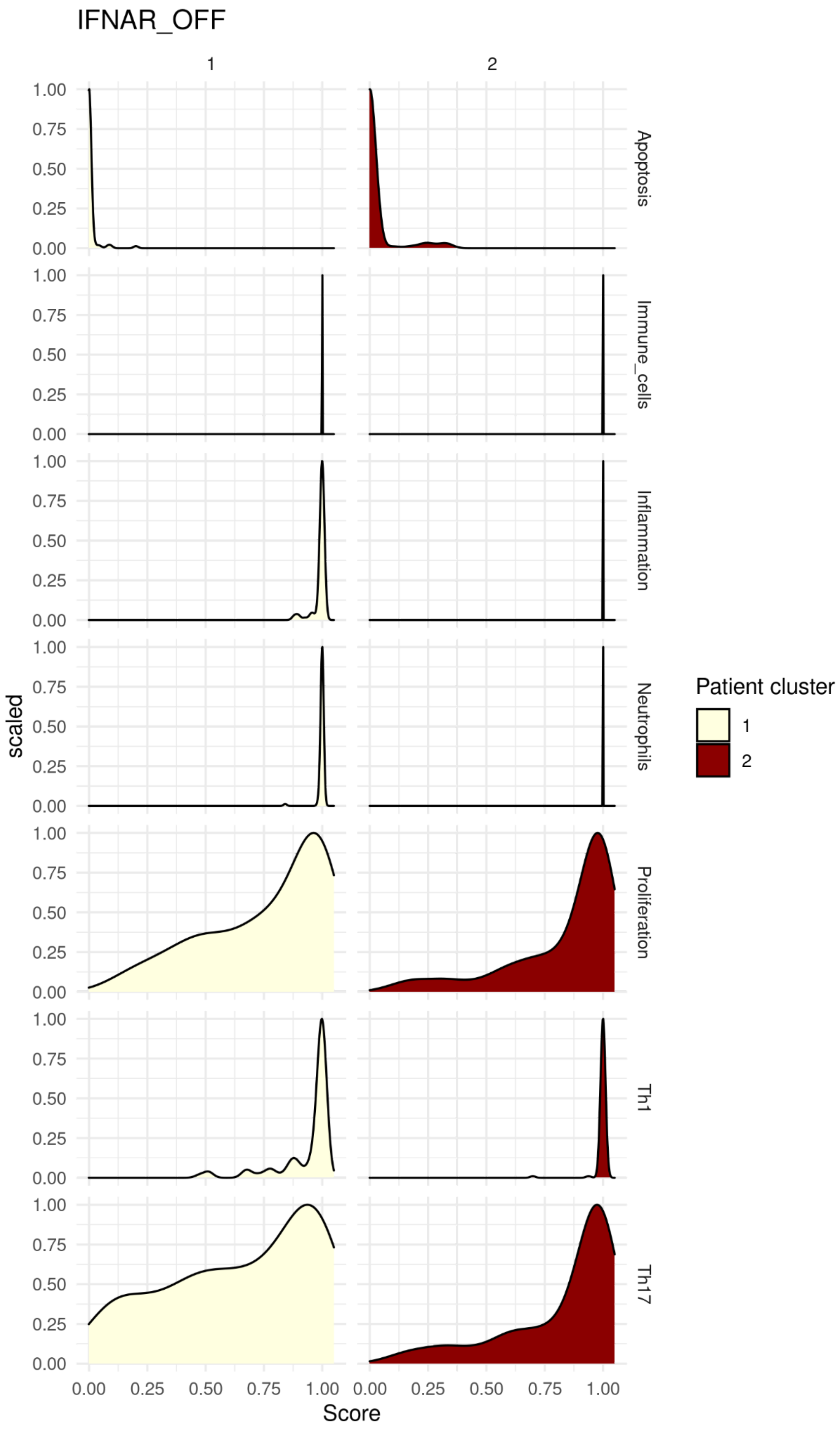
IFNAR_OFF
1
2
Apoptosis
Immune_cells
Inflammation
Neutrophils
Proliferation
Th1
Th17
scaled
Score
0.00
0.25
0.50
0.75
1.00
Patient cluster
1
2

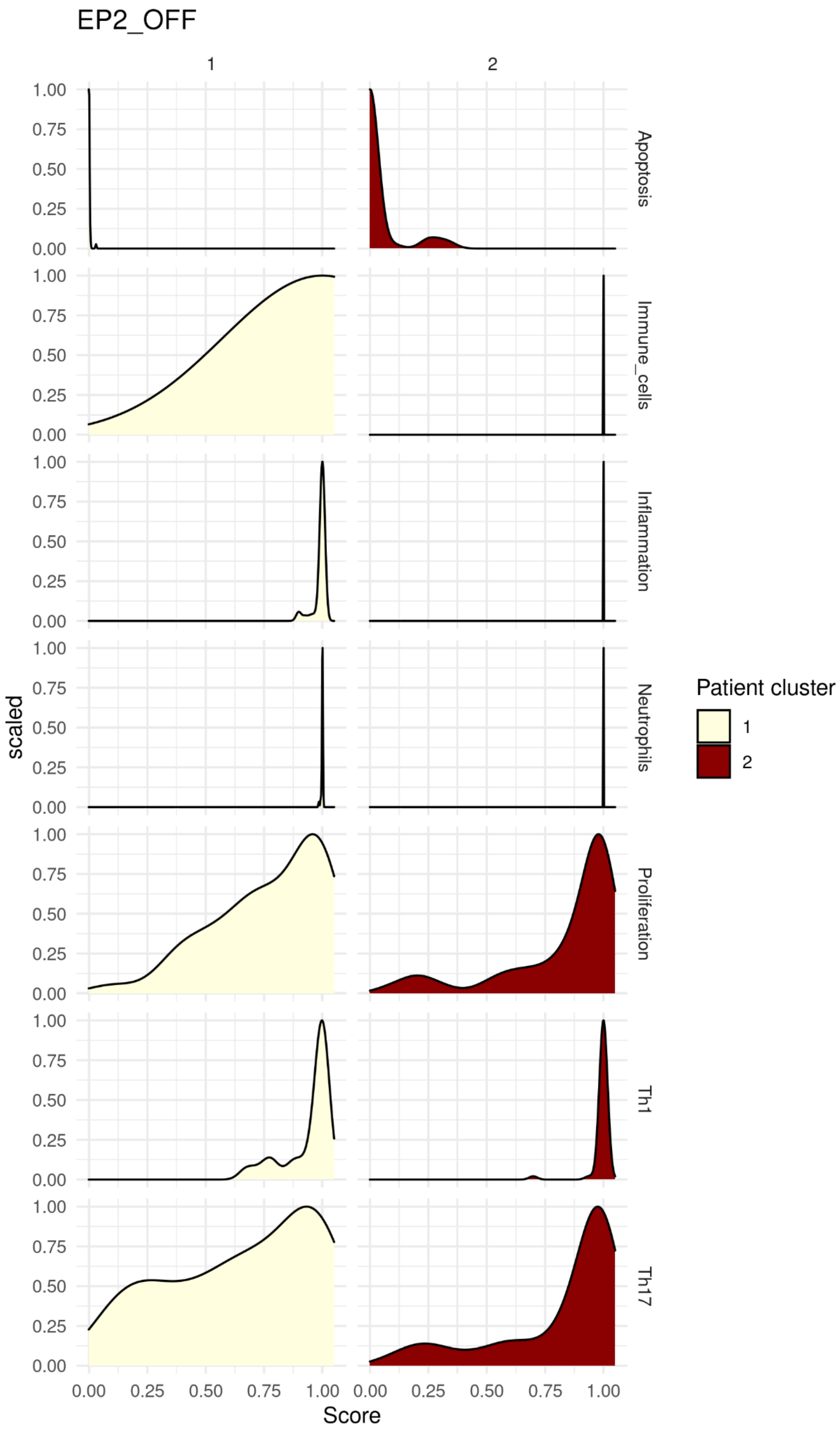

EP2_OFF
1
2
Apoptosis
Immune_cells
Inflammation
Neutrophils
Proliferation
Th1
Th17
scaled
1.00
0.75
0.50
0.25
0.00
Score
0.00 0.25 0.50 0.75 1.00
Patient cluster
1
2

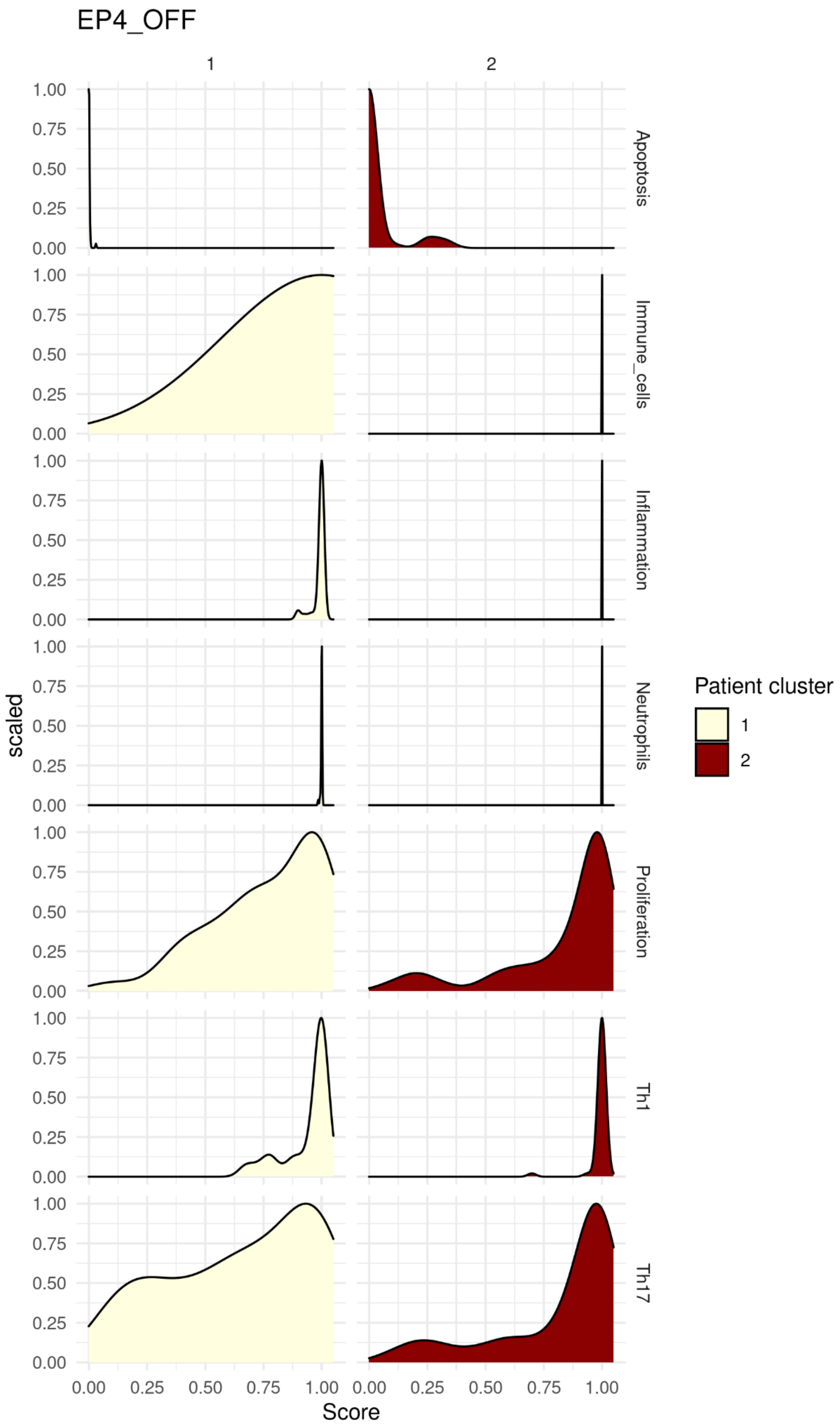

EP4_OFF
1
2
Apoptosis
Immune_cells
Inflammation
Neutrophils
Proliferation
Th1
Th17
scaled
1.00
0.75
0.50
0.25
0.00
0.00 0.25 0.50 0.75 1.00
Score
Patient cluster
1
2

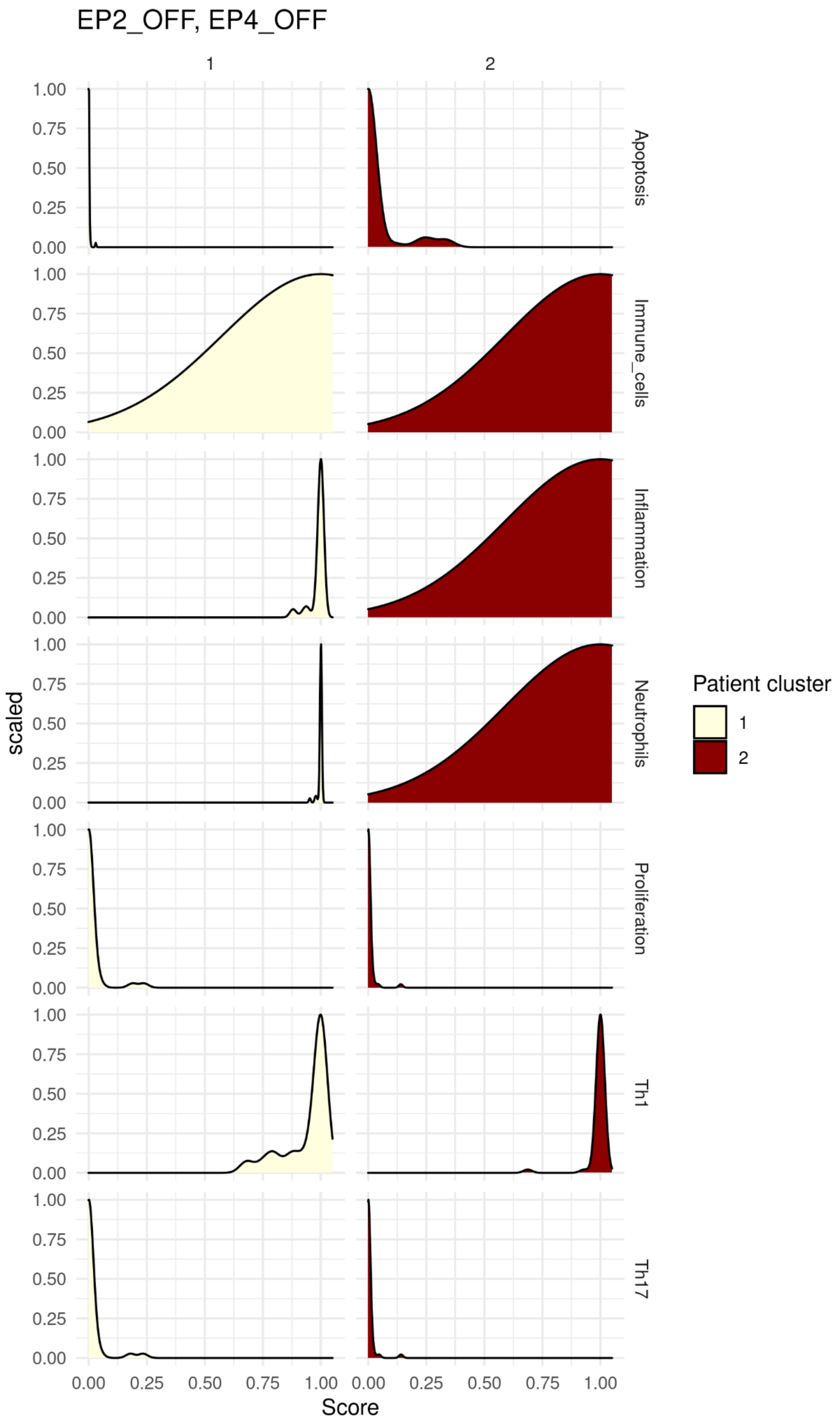

EP2_OFF, EP4_OFF
1
2
Apoptosis
Immune_cells
Inflammation
Neutrophils
Proliferation
Th1
Th17
scaled
Score
0.00
0.25
0.50
0.75
1.00
Patient cluster
1
2

**Supplementary Table 1.** Outline of the nine studies that were employed. The studies' GSE ids and publication references (PMIDs) are presented with the total psoriasis samples included in the study and the number of samples that were used in this project from the respective datasets. The reason for why specific samples were discarded is also explained.

| Publication | Studies | Total samples | Used samples | Reason for sample removal |
|---|---|---|---|---|
| PMID:21850022 | GSE41745 | 6 | 6 | |
| PMID:26251673 | GSE67785 | 28 | 28 | |
| PMID:24441097 | GSE54456 | 174 | 174 | |
| PMID:29031600 | GSE83645 | 25 | 25 | |
| PMID:25723451 | GSE63979 | 42 | 42 | |
| PMID:30054515 | GSE117405 | 28 | 28 | |
| PMID:24909886 | GSE47944 | 84 | 21 | 63 samples discarded because of AhR-antagonist, DMSO or AhR-agonist treatments |
| PMID:27793094 | GSE74697 | 52 | 34 | 18 samples discarded because of adalimumab treatment |
| PMID:29273799 | GSE107871 | 24 | 12 | 12 samples discarded because of cell cultivation differences |

**Supplementary Table 3.** An overview of the 66 marker genes that the logical model includes after the severity alteration. The respective phenotype the gene has is indicated, and whether the gene is up- or downregulated in cluster 2 compared to cluster 1 is also outlined.

| Marker genes | Phenotype | DEG status |
|---|---|---|
| BAD | Apoptosis | |
| CASP8 | Apoptosis | |
| FOXO3 | Apoptosis | |
| IFI6 | Survival | Upregulated |
| IL15 | Survival | |
| CFLAR | Survival | |
| BCL2 | Survival | |

| | | |
|---|---|---|
| CCND1 | Survival | |
| CREB1 | Proliferation | |
| IL8 | Proliferation | |
| IL19 | Proliferation | |
| PGE2 | Proliferation | |
| HETE12 | Proliferation | |
| PPARD | Proliferation | |
| HBEGF | Proliferation | |
| KRT6 | Proliferation | Upregulated |
| BIRC3 | Proliferation | |
| DEFB3 | Proliferation | |
| TIMP1 | Proliferation | |
| MYC | Proliferation | |
| FLG | Differentiation | Downregulated |
| KRT1 | Differentiation | |
| CALML5 | Differentiation | |
| SIRT1 | Differentiation | |
| CDKN1A | Differentiation | |
| WIF1 | Differentiation | Downregulated |
| LOR | Differentiation | Downregulated |
| IL6 | Immune cells | |
| CCL7 | Immune cells | |
| CXCL1 | Immune cells | Upregulated |
| CXCL2 | Immune cells | |
| CXCL5 | Immune cells | |
| LTB4 | Immune cells | |
| CSF3 | Immune cells | |
| ICAM1 | Immune cells | |

| | | |
|---|---|---|
| NOS2 | Immune cells | Upregulated |
| CXCL11 | Immune cells | |
| ISG15 | Immune cells | |
| IFNB | Immune cells | |
| CCL3 | Immune cells | |
| S100A7A | Immune cells | Upregulated |
| IFNA | Immune cells | |
| S100A7 | Inflammation | |
| S100A8 | Inflammation | |
| S100A9 | Inflammation | |
| DEFB4A | Inflammation | Upregulated |
| EP2_g | Inflammation | |
| EP4_g | Inflammation | |
| OAS2 | Inflammation | Upregulated |
| S100A12 | Inflammation | Upregulated |
| LCN2 | Inflammation | Upregulated |
| OASL | Inflammation | Upregulated |
| MX1 | Inflammation | Upregulated |
| RSAD2 | Inflammation | Upregulated |
| SERPINB3_4 | Inflammation | Upregulated |
| CXCL3 | Th1 | |
| IFNG | Th1 | |
| IL12 | Th1 | |
| IL23 | Th17/22 | |
| IL36A | Th17/22 | Upregulated |
| IL36G | Th17/22 | Upregulated |
| IL1B | Th17/22 | Upregulated |
| TNFa | Th17/22 | |

| | |
|---|---|
| CCL2 | Th17/22 |
| CCL5 | Th17/22 |
| CCL20 | Th17/22 |

**Supplementary Table 4.** Overview of the validation of the "IL1B and IL36 pathway." Gene expressions/activities from literature during KC IL36 stimulus are presented. ON signifies an upregulated expression after this experimental stimulus, whereas OFF signifies downregulation following such stimulus. Respective gene activities in the severity model when performing the "IL1B and IL36 pathway" perturbation are also presented; the activities are presented for the pathway activation without or with a simultaneous activation of prostaglandin signaling via EP2 and EP4.

| Marker genes | Experimental activity | Model activity following IL1B and IL36 pathway stimulation | Model activity with simultaneous inclusion of EP2 and EP4 stimulation |
|---|---|---|---|
| CXCL1 | ON (PMID:24829417) | ON | ON |
| CCL3 | ON (PMID:24829417) | OFF | ON |
| CCL5 | ON (PMID:24829417) | OFF | ON |
| CCL20 | ON (PMID:24829417) | ON | ON |
| CCL2 | ON (PMID:24829417) | OFF | ON |
| IL36A | ON (PMID:27783881, PMID:31284527) | OFF | ON |
| IL36G | ON (PMID:27783881, PMID:29142248) | OFF | ON |
| S100A8 | ON (PMID:31284527) | ON | ON |
| DEFB3 | ON (PMID:31284527) | ON | ON |
| S100A7 | ON (PMID:31284527) | ON | ON |
| TNFa | ON (PMID:31284527) | ON | ON |
| IL6 | ON (PMID:31284527) | ON | ON |
| LCN2 | ON (PMID:31284527) | OFF | ON |
| S100A7A | ON (PMID:31284527, PMID:29142248) | ON | ON |
| CXCL3 | ON (PMID:31284527) | ON | ON |
| DEFB4A | ON (PMID:29142248) | OFF | ON |
| FLG | OFF (PMID:29142248) | OFF | OFF |
| LOR | OFF (PMID:29142248) | OFF | OFF |
| IL1B | ON (PMID:31539532) | OFF | ON |

**Supplementary Table 5.** Presentation of the gene expressions/activities of different marker genes retrieved from scientific publications that describe IL1B KC stimuli. The respective activities in the severity model when performing a "IL36 and IL1B pathway" simulation is indicated, both without and with a simultaneous activation of signaling via EP2 and EP4.

| Marker genes | Experimental activity | Model activity following IL36 and IL1B pathway stimulation | Model activity with simultaneous signaling via EP2 and EP4 |
|---|---|---|---|
| CCL20 | ON (PMID:29434599) | ON | ON |
| S100A7A | ON (PMID:29434599) | ON | ON |
| S100A7 | ON (PMID:29434599) | ON | ON |
| IL1B | ON (PMID:29434599, PMID:33584721) | OFF | ON |
| IL23A | ON (PMID:29434599) | OFF | OFF |
| IL36A | ON (PMID:29434599) | OFF | OFF |
| IL36G | ON (PMID:29434599) | OFF | OFF |
| CXCL1 | ON (PMID:29434599) | ON | ON |
| CXCL2 | ON (PMID:29434599) | OFF | ON |
| CXCL5 | ON (PMID:29434599) | ON | ON |
| TNFa | ON (PMID:33584721) | ON | ON |
| CCL5 | ON (PMID:19166933) | OFF | ON |
| CXCL11 | ON (PMID:19166933) | OFF | ON |
| CCL2 | ON (PMID:19166933) | OFF | ON |
| IL6 | ON (PMID:19166933) | ON | ON |
| ICAM1 | ON (PMID:7913702, PMID:25238321) | OFF | ON |
| S100A12 | ON (PMID:21316034) | ON | ON |
| NOS2 | ON (PMID:15275864) | OFF | ON |
| KRT6 | ON (PMID:11180011) | ON | ON |
| BIRC3 | ON | ON | ON |

**Supplementary Table 6.** Overview of the activity of the marker genes in the severity model that literature has revealed as relevant during IFN alpha or beta signaling in KCs. ON signifies that the gene is produced following the stimulation of IFN alpha/beta. The genes' stable state activities when performing an *in silico* simulation with the "IFNAR" state are also depicted; that is when IFN alpha/beta signaling is activated in the severity model.

| Marker gene | Experimental activity | Model activity |
|---|---|---|
| IL6 | ON (PMID:17928888) | ON |
| ICAM1 | ON (PMID:17928888) | ON |
| ISG15 | ON (PMID:18648529) | ON |
| IFI6 | ON (PMID:18648529) | ON |
| RSAD2 | ON (PMID:18648529) | ON |
| MX1 | ON (PMID:18648529) | ON |
| OAS2 | ON (PMID:18648529) | ON |
| OASL | ON (PMID:18648529) | ON |
| CXCL11 | ON (PMID:28472186) | ON |
| CCL5 | ON (PMID:28472186) | ON |
| IFNB | ON (PMID:30936491) | OFF |
| IFNA | ON (PMID:30936491) | ON |
| IFNG | ON (PMID:30936491) | ON |

**Supplementary Table 7.** A representation of the expressions/activities of some marker genes that were incorporated in the severity model for the IL6 pathway. "ON" refers to that the scientific publication revealed induced upregulation of these marker genes after IL6 stimuli. The respective activities in the severity model for these marker genes for the "IL6" perturbation are also given.

| Marker nodes | Experimental activity | Model activity |
|---|---|---|
| TIMP1 | ON (PMID:27297362) | ON |
| MYC | ON (PMID:21159631) | ON |
| IL15 | ON (PMID:21769475) | ON |
| SERPINB3_4 | ON (PMID:27637160) | ON |

**Supplementary Table 8.** An overview of the model validation performed with the "Psoriasis state" perturbation. The stable state gene activities from the corresponding "ALL" situation in Tsirvouli et al. is listed for the readout genes that the severity model shares with the base model. Additionally, for the readout genes that the severity model did not share with the base model, the gene's level of activity is retrieved from additional publications that uncover the implication of these genes when it comes to psoriasis pathology. ON implies that the gene is over-expressed, while OFF implies that it is under-expressed in psoriasis. Lastly, the activity of these readout genes in the severity model following the *in silico* "*Psoriasis state*" perturbation is presented.

| Markers | Tsirvouli et al. "ALL condition" | Experimental activity | Model activity |
|---|---|---|---|
| BAD | OFF | | OFF |
| CASP8 | OFF | | OFF |
| FOXO3 | OFF | | OFF |
| IFI6 | | ON (PMID:20377629, PMID:31539532) | ON |
| IL15 | | ON (PMID:10925312, PMID:16329645, PMID:24086722) | ON |
| CFLAR | ON | | ON |
| BCL2 | ON | | ON |
| CCND1 | ON | | ON |
| CREB1 | ON | | ON |
| IL8 | ON | | ON |
| IL19 | ON | | ON |
| PGE2 | ON | | ON |
| HETE12 | ON | | ON |
| PPARD | ON | | ON |
| HBEGF | ON | | ON |
| KRT6 | | ON (PMID:11180011, PMID:26849645) | ON |
| BIRC3 | | ON (PMID:24372854) | ON |

| | | | |
|---|---|---|---|
| DEFB3 | | ON<br>(PMID:30320469, PMID:34480893) | ON |
| TIMP1 | | ON<br>(PMID:11380613, PMID:14523226) | ON |
| MYC | | ON<br>(PMID:1688596,<br>PMID:26046687,<br>PMID:17673382) | ON |
| FLG | OFF | | OFF |
| KRT1 | OFF | | OFF |
| CALML5 | OFF | | OFF |
| SIRT1 | OFF | | OFF |
| CDKN1A | ON | | ON |
| WIF1 | | OFF<br>(PMID:31987884, PMID:20376066) | OFF |
| LOR | | OFF<br>(PMID:33458806, PMID:15598222,<br>PMID:15187321) | OFF |
| IL6 | ON | | ON |
| CCL7 | ON | | ON |
| CXCL1 | ON | | ON |
| CXCL2 | ON | | ON |
| CXCL5 | ON | | ON |
| LTB4 | ON | | ON |
| CSF3 | ON | | ON |
| ICAM1 | | ON<br>(PMID:19022587, PMID:34414905) | ON |
| NOS2 | | ON<br>(PMID:15275864, PMID:34909719) | ON |
| CXCL11 | | ON<br>(PMID:28472186, PMID:18684158) | ON |
| ISG15 | | ON<br>(PMID:24260178, PMID:25809693) | ON |

| | | | |
|---|---|---|---|
| IFNB | | ON<br>(PMID:31293591, PMID:31010609) | ON |
| CCL3 | | ON<br>(PMID:23223135, PMID:30760013) | ON |
| S100A7A | | ON<br>(PMID:20220767, PMID:26055798) | ON |
| IFNA | | ON<br>(PMID:31293591) | ON |
| S100A7 | ON | | ON |
| S100A8 | ON | | ON |
| S100A9 | ON | | ON |
| DEFB4A | ON | | ON |
| EP2_g | ON | | ON |
| EP4_g | ON | | ON |
| OAS2 | | ON<br>(PMID:32849499, PMID:22071477) | ON |
| S100A12 | | ON<br>(PMID:20220767, PMID:26333514) | ON |
| LCN2 | | ON<br>(PMID:30224457, PMID:26849645) | ON |
| OASL | | ON<br>(PMID:18648529, PMID:31539532) | ON |
| MX1 | | ON<br>(PMID:32648291, PMID:19399181) | ON |
| RSAD2 | | ON<br>(PMID:25809693, PMID:32648291) | ON |
| SERPINB3_4 | | ON<br>(PMID:30054515, PMID:22071477,<br>PMID:32962824) | ON |
| CXCL3 | ON | | ON |
| IFNG | ON | | ON |
| IL12 | ON | | ON |
| IL23 | ON | | ON |

| | | | |
|---|---|---|---|
| IL36A | ON | | ON |
| IL36G | ON | | ON |
| IL1B | ON | | ON |
| TNFa | ON | | ON |
| CCL2 | ON | | ON |
| CCL5 | ON | | ON |
| CCL20 | ON | | ON |
